# Epigenome-informed prioritization of bivalent chromatin SNPs enhances genomic prediction robustness: a proof-of-concept study in Pacific white shrimp (*Litopenaeus vannamei*)

**DOI:** 10.64898/2026.02.14.705940

**Authors:** Jiale Shi, Zhifang Lu, Mingyi Sui, Miaomiao Yin, Ming Mu, Dongdong Zhang, Zhenmin Bao, Jingjie Hu, Qifan Zeng, Zhi Ye

**Affiliations:** Key Laboratory of Tropical Aquatic Germplasm of Hainan Province, Sanya Oceanographic Institution, Ocean University of China, Sanya 572024, China; MOE Key Laboratory of Marine Genetics and Breeding, College of Marine Life Sciences, Ocean University of China, Qingdao 266003, China; School of Breeding and Multiplication (Sanya Institute of Breeding and Multiplication), School of Marine Biology and Fisheries, Hainan University, Sanya 572025, China

**Keywords:** Genomic selection, Histone modifications, Epigenetics, CUT&Tag, Functional annotation

## Abstract

**Background:** Genomic selection (GS) has revolutionized animal breeding, spanning livestock sectors such as pigs and cattle to aquatic species like fish and shrimp. However, its broader application across these industries is often constrained by high genotyping costs and reduced predictive reliability across divergent populations or generations. Developing cost-effective, biologically informed genotyping strategies to overcome these limitations remains a critical goal in animal agriculture. Epigenetic annotations, particularly histone modifications, provide direct functional insights into regulatory elements underlying complex trait variation and represent a promising but underexplored resource for marker prioritization.

**Results:** Here, using the Pacific white shrimp (*Litopenaeus vannamei*) as a model organism, we conducted a proof-of-concept study integrating resequencing and phenotypic data from 972 individuals. We generated high-resolution epigenomic maps by profiling four histone marks (H3K4me1, H3K4me3, H3K27me3, and H3K27ac) across multiple embryonic stages and adult muscle tissue using CUT&Tag. These functional annotations were then leveraged to prioritize single nucleotide polymorphism (SNP) subsets for genomic prediction. Among the tested strategies, SNPs located in the muscle-specific bivalent promoter/enhancer (E6) state—characterized by the co-occurrence of active and repressive marks—consistently maximized prediction accuracy under the BayesA model. Notably, even at a moderate density (15k), E6-derived SNPs achieved prediction accuracies exceeding those obtained using substantially larger genome-wide SNP sets. Most importantly, in a challenging cross-population validation using an independent strain, the E6-derived SNP subset significantly improved prediction accuracy by 47.6% (increasing from 0.21 ± 0.05 to 0.31 ± 0.04, *p* < 0.05) compared to random subsets at equivalent density.

**Conclusions:** These results demonstrate that epigenetic annotation–guided SNP prioritization provides a biologically informed and cost-effective strategy to enhance genomic prediction accuracy and stability. This framework is broadly transferable across species and offers a practical strategy for designing low-density genotyping panels that reduce costs while maintaining reliable selection outcomes in large-scale breeding programs.

## 1. Introduction

Genomic selection (GS) has fundamentally transformed animal breeding by accelerating genetic gain for complex traits, spanning from established livestock sectors such as pigs and cattle [1,2] to emerging aquaculture species including fish, shrimp, and shellfish [3]. By estimating breeding values using genome-wide molecular markers, GS circumvents the limitations of traditional pedigree-based selection. However, a critical bottleneck remains: the predictive accuracy of GS models often declines rapidly when applied across genetically divergent populations or across generations, largely due to the decay of linkage disequilibrium (LD) between markers and causal variants [4,5]. While whole-genome sequencing (WGS) data theoretically captures all causal mutations, studies have indicated that simply increasing marker density does not guarantee improved prediction accuracy [3,6,7]. For instance, Zhuang et al. (2023) demonstrated in a commercial pig population that utilizing full WGS data did not significantly outperform single nucleotide polymorphism (SNP) chips, primarily due to the accumulation of "noisy" variants in non-functional genomic regions that dilute predictive signals [8].

To overcome these limitations, a paradigm shift is occurring from purely statistical modeling toward integrating "biological priors"—functional genomic annotations that pinpoint regulatory elements [9]. Recent high-impact studies across diverse species have demonstrated that prioritizing variants within functional regions (e.g., promoters and enhancers) significantly enhances the robustness of genomic prediction [5,6,10]. In cattle, Xiang et al. (2025) recently integrated extensive multi-omics annotations to develop a functional importance score, which improved the genomic prediction accuracy for climate resilience traits by approximately 11% across diverse populations [11]. Similarly, Ma et al. (2025) developed a deep learning-based framework (DeepAnnotation) for pigs, showing that integrating functional annotations can effectively filter out noise and improve phenotype prediction [12]. These findings suggest that identifying functional regulatory variants is key to breaking the LD dependency and achieving stable predictions across diverse genetic backgrounds. However, despite these encouraging breakthroughs, the integration of functional annotation into genomic prediction is still considered to be in its developmental stages [13]. Current applications are largely restricted to a few model livestock species and specific traits, and the optimal strategies for variant prioritization remain debated [12]. To establish stable, reference-grade protocols that meet diverse breeding demands, it is essential to validate these functional genomics strategies across a wider spectrum of species and complex traits.

Among the various layers of functional information, epigenetic modifications—particularly histone post-translational modifications (PTMs)—serve as dynamic indicators of the genomic regulatory landscape [14–16]. Specific histone marks act as fundamental signatures for defining cis-regulatory elements (CREs); for instance, trimethylation of histone H3 at lysine 4 (H3K4me3) typically marks active promoters, whereas acetylation at lysine 27 (H3K27ac) distinguishes active enhancers from inactive ones. However, relying on individual marks provides only a partial view of regulatory complexity. By integrating profiles of multiple histone modifications (e.g., H3K4me1, H3K4me3, H3K27ac, and H3K27me3), the genome can be segmented into distinct "chromatin states" (e.g., active, repressed, or poised regions), providing a high-resolution map of functional elements [14,17,18]. These chromatin states serve as powerful "biological priors" for genomic prediction, enabling the prioritization of SNPs located in biologically active regulatory regions while filtering out noise from non-functional sequences. Consequently, leveraging these multi-mark chromatin states offers a mechanism-based strategy to refine SNP selection and potentially enhance the predictive power of GS models.

In the aquaculture sector, concerted international initiatives like AQUA-FAANG (Functional annotation of animal genomes consortium) have begun to characterize functional regulatory landscapes for major European finfish species [19], complemented by similar FAANG-associated efforts in the United States targeting key species such as rainbow trout [14]. Emerging evidence from these projects indicates that functional genomic annotations can successfully assist in pinpointing causative variants underlying complex traits, such as disease resistance in sea bass [19] and rainbow trout [14]. However, despite this progress in map generation, the empirical application of functional genomic annotations into GS models remains unreported in aquaculture. Furthermore, compared to finfish, functional genomic resources for economically critical crustacean species— particularly the Pacific white shrimp (*Litopenaeus vannamei*)—are severely lacking. While high-quality reference genomes for shrimp are now available, the comprehensive epigenetic maps required to filter "noisy" variants and guide robust genomic prediction are virtually nonexistent, limiting the translation of functional genomics into practical shrimp breeding.

To address this gap, this study presents a proof-of-concept framework for epigenetically informed genomic selection in *L. vannamei*. We generated high-resolution chromatin maps using CUT&Tag (Cleavage under targets and tagmentation) for four key histone modifications (H3K4me1, H3K4me3, H3K27me3, and H3K27ac) across embryonic development and adult tissues. By integrating these functional annotations with whole-genome resequencing data from a breeding population, we systematically evaluated the predictive efficiency of different chromatin states. We specifically hypothesized that variants located in bivalent promoter/enhancer regions (E6 state) would offer superior predictive robustness compared to random genome-wide markers. Our findings demonstrate that prioritizing these functional SNPs significantly improves cross-population prediction accuracy, offering a cost-effective and biologically grounded strategy for precision breeding in aquaculture.

## 2. Material and methods

### 2.1 Breeding populations

The three breeding populations analyzed in this study were derived from ongoing selective breeding programs conducted in our laboratory. Three domesticated breeding strains of *L. vannamei* maintained at a commercial hatchery (Hairen Aquatic Seed Industry Technology Co., Ltd., Hebei, China) were included. Each strain was produced from approximately 50 breeding pairs of broodstock through mass spawning, generating large mixed-family larval cohorts of approximately 5,000,000 offspring per strain. Individuals used in this study were randomly sampled from these cohorts for genotyping and phenotyping until they reach the desired age under rearing conditions described below. Due to the mass-spawning procedure, physical pedigree records (i.e., full- and half-sib family structure) were not available. Therefore, realized genetic relationships among individuals were estimated using a genomic relationship matrix derived from genome-wide SNP markers. In addition, a 110-day-old Hawaiian Primo (PLM) strain population from Haimao Seed & Breeding Technology Group Co., Ltd. (Guangdong, China) was included as an external, distantly related population for cross-population evaluation. This population originated from a single breeding family producing approximately 200,000 offspring, from which individuals were randomly sampled.

All shrimp were reared in controlled indoor recirculating aquaculture systems prior to sampling. Seawater was subjected to a standardized treatment process consisting of ozonation, ultraviolet sterilization, and fine mechanical filtration (1–5 μm). Each tank (effective volume 80–100 m³) was stocked at a density of 20–25 individuals/m³. During the rearing period, water temperature was maintained at 25–28°C, salinity at 28–31‰, and pH at 7.8–8.1. Dissolved oxygen levels were kept above 6 mg/L, while concentrations of ammonia-N and nitrite-N were controlled below 3 mg/L and 0.05 mg/L, respectively. To ensure stable environmental conditions, one-third to one-half of the tank volume was renewed daily. Shrimp were fed a high-protein formulated diet four times per day at a feeding rate of 12–15% of total biomass.

#### 2.1.1 Phenotypic statistics

Body length and body weight were obtained using a digital caliper (± 0.02 mm) and an electronic scale (± 0.01 g), respectively. Summary statistics for each population are presented in **Table 1**.

**Table 1.** Phenotypic statistics of growth traits in *L. vannamei*.

| Group | Sample size | Length (mm)<br>(Mean ± SEM) | Weight (g)<br>(Mean ± SEM) |
| --- | --- | --- | --- |
| 1 | 313 | 221.14 ± 1.01 | 66.79 ± 0.73 |
| 2 | 331 | 200.18 ± 0.68 | 49.27 ± 0.43 |
| 3 | 328 | 169.95 ± 0.64 | 31.18 ± 0.30 |
| PLM | 73 | 133.82 ± 0.10 | 18.90 ± 0.44 |

To assess phenotypic distributions, boxplots of the body length and body weight were generated using the R package ggplot2 [20]. Normality of body length and body weight was tested with IBM SPSS Statistics v25.0. The correlation between body weight and body length was evaluated by linear regression, and coefficients of determination (R²) were calculated in Microsoft Excel 2021.

#### 2.1.2 Resequencing, Genotyping, SNP discovery and quality control

Genomic DNA was extracted from muscle tissue using the Tiangen Genomic DNA Extraction Kit (Beijing, China). Sequencing libraries were prepared with the VAHTS Universal DNA Library Prep Kit for Illumina V3® (Vazyme, China) and sequenced on the Illumina NovaSeq 6000 platform (Illumina, USA) to generate 150 bp paired-end reads, targeting approximately 2 Gb of sequencing data per individual.

Raw sequencing reads were quality assessed and filtered using the NGS QC Toolkit v2.3 [21]. Reads containing adapter contamination, >10% ambiguous bases (N), or >40% bases with Phred quality < 20 were removed. High-quality reads were aligned to the chromosome-level *L. vannamei* reference genome available through the Aquaculture Molecular Breeding Platform (AMBP; http://mgb.qnlm.ac/) [22] using BWA-MEM v0.7.17 [23]. PCR duplicates were removed using the MarkDuplicates tool in GATK v4.0 [24], and alignment files were sorted and indexed with SAMtools v1.17 [25].

Variants were called using the GATK v4.0 [24] best-practice pipeline. The *HaplotypeCaller* was run in GVCF mode for each sample with default parameters, followed by CombineGVCFs and GenotypeGVCFs to produce a joint VCF (Variant call format) file containing all individuals. To ensure variant accuracy, only biallelic SNPs were retained using the GATK SelectVariants tool (flag –select-type-to-include SNP). Variants were filtered using the VariantFiltration function in GATK v4.0, and loci with Quality by Depth (QD) < 2.0, Fisher Strand Bias (FS) > 60.0, Mapping Quality (MQ) < 40.0, Mapping Quality Rank Sum (MQRankSum) < -12.5, Read Position Rank Sum (ReadPosRankSum) < -8.0, or Strand Odds Ratio (SOR) > 3.0 were excluded from downstream analyses.

Genotypes were subsequently imputed using GLIMPSE v2.0.0 [26] with our in-house haplotype reference panel. SNP quality control was performed using Plink v1.9 [27]: SNPs with minor allele frequency (MAF) < 0.05 were excluded, yielding 2,978,750 SNPs. Hardy–Weinberg equilibrium (HWE) was tested (--hwe 0.0001), and loci with significant deviations (*p* < 0.0001) were removed, resulting in 2,609,549 high-quality SNPs for downstream analysis. Sample- and locus-level missingness was assessed using PLINK (--missing), and variant statistics were visualized in R using ggplot2 [20].

#### 2.1.3 Population structure and individual genetic relatedness

Population structure was assessed using both principal component analysis (PCA) and model-based clustering. PCA was performed with Plink v1.9 (--indep-pairwise 50 10 0.5; --pca 10) to remove linkage disequilibrium and extract the top ten principal components (PCs), which were used to visualize genetic stratification among populations. Model-based population structure analysis was conducted using ADMIXTURE v1.3.0 [28] with the number of assumed ancestral clusters (K) ranging from 2 to 30. Because the cross-validation (CV) error decreased gradually with increasing K without showing a clear minimum (Fig. S2A), the choice of K was guided by the stability of ancestry patterns across replicate runs, the consistency with PCA clustering, and the known breeding histories of the strains. Based on these criteria, K = 3 was selected as the most parsimonious representation of the major population structure. Kinship among individuals was estimated using GEMMA v0.98.5 [29] to construct the relatedness matrix. Visualization of PCA, ADMIXTURE ancestry coefficients, and kinship matrices was performed using the R packages ggplot2 [20] and ComplexHeatmap [30]. SNP density plots were generated with CMplot [31].

#### 2.1.4 Estimation of genetic parameters based on SNP markers

The narrow-sense heritability (h²) of each trait was estimated using genome-wide SNP markers under a genomic linear mixed model (LMM) implemented in GCTA v1.94.3 [32]. The LMM was specified as:

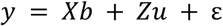

where *y* is the vector of phenotypic observations, *X* is the design matrix for fixed effects, and *b* is the vector of fixed-effect coefficients. Because the three breeding populations were reared separately and environmental records were unavailable, population identity was included as the fixed effect to prevent confounding and inflation of additive genetic variance due to between-population divergence. *Z* is the incidence matrix relating records to individuals, and *u* ∼ 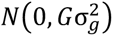 represents the additive genetic effects, with *G* being the genomic relationship matrix (GRM) constructed from SNP genotypes using the command *--make-grm*. The residual term is denoted as 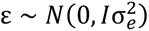.

Variance components 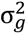 (additive genetic variance) and 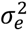 (residual variance) were estimated using restricted maximum likelihood (REML) with the parameters *--reml* and *--reml-alg 0*. The REML algorithm iteratively maximizes the restricted log-likelihood function to obtain optimal estimates of the variance components. The heritability of each trait was then calculated as:

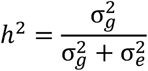

Because population and environmental effects were confounded, this formulation estimates the within-population heritability, reflecting additive genetic variance after removing population mean differences. This approach ensures that the estimated h² values represent true intra-population genetic variation, rather than inflated values driven by between-population divergence.

### 2.2 Genome-wide histone modification information acquisition

#### 2.2.1 CUT&Tag sample description

The CUT&Tag samples used in this study comprised specimens from both embryonic developmental stages and adult muscle tissue. Embryonic samples covered seven stages—blastula (blas), gastrula (gast), limb bud embryo (lbe), larva in membrane (lim), nauplius I (N1), nauplius III (N3), and nauplius VI (N6)—which were obtained from our previously published work [33]. In addition, CUT&Tag profiling was conducted on adult muscle tissue collected from healthy shrimp, which were aseptically dissected for histone modification analysis.

To extend the developmental coverage, additional CUT&Tag datasets were generated for four later embryonic stages (zoea I, zoea III, mysis I, and mysis III). These newly generated datasets have been submitted to the NCBI BioProject database and will be made publicly available upon publication of this article.

#### 2.2.2 CUT&Tag experimental procedure and data analysis

Two independent biological replicates were performed for each histone modification under identical experimental conditions to ensure reliability and reproducibility. Approximately 200 embryos were collected at each developmental stage per replicate, and ∼50,000 nuclei were used for each CUT&Tag experiment. For adult samples, approximately 100 mg of dorsal muscle tissue was aseptically dissected from healthy shrimp, and nuclei were isolated following the protocol described in Qi et al. (2025). The CUT&Tag experiments were performed following the principles described by Kaya-Okur et al. (2019). Libraries were prepared using the Illumina Hyperactive Universal CUT&Tag Kit (Vazyme, China; Cat. #TD903), with minor modifications for *L. vannamei* embryos and muscle tissues. The experimental workflow was consistent with that described in Shi et al. (2025b) for the embryonic samples and Qi et al. (2025) for the muscle tissue. Information on the four histone modification antibodies (H3K4me1, H3K4me3, H3K27me3, and H3K27ac) and the control IgG is provided in **Table S1**. Sequencing was conducted on an Illumina NovaSeq 6000 platform (Novogene, Tianjin, China) with an average depth of ∼1 Gb per sample.

CUT&Tag data analysis was performed using the standardized pipeline established in our previous study [33]. Specifically, raw reads were quality-controlled using NGS QC Toolkit v2.3 [21] by removing adapter sequences, reads containing >10% ambiguous bases (N), or reads with >50% of bases having Phred quality scores ≤20 (summary in **Table S2**). Clean reads were then aligned to the *L. vannamei* reference genome using BWA v0.7.17 [22,23]. Abnormally paired fragments were filtered using the *awk* command to retain valid fragments between 100–1000 bp. Peak calling was then conducted using the remaining fragments with MACS2 v2.2.7.1 [34] using the following parameters: macs2 callpeak -t A.bam -c IgG.bam -f BAMPE -g 1.66e9 --keep-dup auto -q 0.05. Valid peaks were annotated using the R packages GenomicFeatures [35] and ChIPseeker [36].

#### 2.2.3 Chromatin state analysis

Integration of multiple histone modification datasets enables the identification of chromatin states such as active enhancers, active transcription start sites, and silencers, thereby providing high-resolution functional annotation of the genome [17]. In this study, CUT&Tag datasets for four histone modifications (H3K4me1, H3K4me3, H3K27me3, and H3K27ac) from 11 embryonic and larval developmental stages, as well as adult muscle tissue of *L. vannamei*, were used for chromatin state identification.

Chromatin state analysis was performed following the standardized pipeline described in our previous study [33] using ChromHMM v1.26 [17]. The *L. vannamei* genome was divided into non-overlapping 200 bp bins, and enrichment signals for each histone modification were binarized into presence/absence states using the BinarizeBam module with parameters -gzip -b 200 -f 0 -g 0 -p 0.0001. Model training and chromatin segmentation were conducted using the LearnModel module with parameters -gzip -d 0.001 -color 129,0,0 -p 20 -i chrhmm. ChromHMM models with varying state numbers (5-15) were compared based on emission probabilities, enrichment patterns, and genomic coverage. Models with fewer than eight states failed to distinguish biologically meaningful features, whereas those with more than eight states introduced redundant categories. Therefore, an eight-state model was selected as the optimal balance between biological interpretability and model complexity. Each chromatin state was subsequently annotated based on combinatorial histone mark patterns, representing distinct functional elements such as active promoters, strong/weak enhancers, repressed or quiescent regions.

### 2.3 Conventional genomic selection (GS) analysis

In conventional GEBV analysis, random SNP sampling was conducted after LD filtering to reduce marker redundancy and ensure approximate independence among loci. High-LD loci were pruned using Plink v1.9 with the parameter --indep-pairwise 50 10 0.5, resulting in 1,081,117 independent SNPs for downstream GS analysis.

To identify the optimal GS model for growth traits in *L. vannamei*, both genomic best linear unbiased prediction (GBLUP) and Bayesian regression approaches were evaluated. For GBLUP, the following linear mixed model was applied:

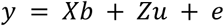

where *y* is the vector of phenotypic records, *b* is the vector of fixed effects including the population group, and *u* is the vector of random additive genetic effects with 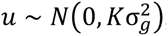. G is the genomic relationship matrix (GRM) which was constructed following VanRaden (2008):

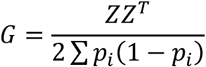

where *Z* is the matrix of centered genotypes and *p*_*i*_ is the allele frequency of the *i* th SNP. Variance components were estimated using the restricted maximum likelihood (REML) algorithm implemented in the R package sommer [37].

In addition to GBLUP, three Bayesian regression models (BayesA, BayesB, and BayesC) were implemented using the BGLR package [38]. The general model was defined as:

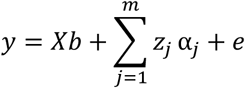

where *z*_*j*_ is the genotype covariate for SNP *j*, and *α*_*j*_ is its effect. The three Bayesian methods differ in their prior assumptions: BayesA assigns marker-specific variances, BayesB introduces a probability (π) that a SNP has zero effect, and BayesC estimates π as a hyperparameter [38,39]. Prediction accuracies derived from each model were compared to determine the best-performing approach.

To assess the effect of marker density on prediction accuracy, SNP subsets ranging from 0.1k to 30k were randomly sampled from the LD-filtered dataset (the 1,081,117 SNPs described above) with the --thin-count parameter in Plink v1.9. For each density, five independent random samplings were performed, and five-fold cross-validation was repeated five times, resulting in 25 replicates per density. To evaluate the impact of training sample size, random subsets of 100, 250, 500, 750, and 972 individuals were used. At a fixed marker density of 10k, loci were randomly sampled five times, and prediction accuracy was evaluated using the BayesA model across 20 rounds of five-fold cross-validation. In five-fold cross-validation, all individuals were randomly divided into five non-overlapping subsets, with one subset used as the validation set and the remaining four as the training set in each iteration until every individual had been used once for validation. Prediction accuracy was defined as the Pearson correlation between predicted GEBVs and observed phenotypes in the validation sets, adjusted by dividing by the square root of trait heritability to account for the influence of finite heritability on predictive performance.

### 2.4 GS analysis based on histone modification-guided SNP filtering

Because it remains unclear which histone modifications (in which tissues) have the greatest impact on genomic selection (GS), we performed a series of stepwise evaluations to comprehensively assess the effects of functional genomic annotations derived from histone modifications on GS performance. Given that histone modifications and chromatin states are dynamically regulated across embryonic development and different tissues, we first used the combined set of histone modification peaks from all tissues to ensure comprehensive coverage of functional genomic regions. Based on this union set, we evaluated the SNP filtering effects of individual histone modifications and compared them with results obtained from muscle tissue alone, which is expected to be more directly related to growth traits.

In addition, because SNPs located in distinct chromatin states may exert different regulatory influences on downstream genes, chromatin state annotations derived from all histone modification data were further applied. Specifically, we evaluated and compared the predictive performance of SNPs located within different chromatin states in embryonic stages and adult muscle tissue separately. The detailed analytical procedures are described below.

#### 2.4.1 Evaluation of GS predictive performance based on single histone modifications

SNPs were filtered based on histone modification peak regions identified from CUT&Tag narrowPeak files. Peak coordinates were extracted using Linux command-line tools (cut and awk), and SNP–peak overlaps were identified with the R package data.table, followed by deduplication. To ensure robustness, only SNPs located within peaks that were consistently detected in both biological replicates were retained. Functional annotation was performed with the R packages GenomicFeatures [35] and ChIPseeker [36], with a *flankDistance* of 5 kb.

From these peak-associated SNPs, five random subsets at different densities were sampled for each histone mark (H3K4me1, H3K4me3, H3K27me3, H3K27ac) and their union set (*All-Histone*) using the --thin-count parameter in Plink v1.9. Parallel subsets were generated from muscle-specific peaks. Genomic prediction was primarily performed using the BayesA model, with predictive performance evaluated by five rounds of five-fold cross-validation. Prediction accuracies were compared with conventional GS (random LD-filtered SNPs), and accuracy was calculated as described in Section 2.3. At equal marker densities, statistical significance was tested using the Wilcoxon rank-sum test (\**p* < 0.05, \*\**p* < 0.01, \*\*\**p* < 0.001), and results were visualized using ggplot2 [20].

#### 2.4.2 Evaluation of GS predictive performance based on chromatin state

Chromatin state profiles for each embryonic stage and adult muscle tissue were generated using the eight-state ChromHMM model described in Section 2.2.3. SNPs passing quality control (prior to LD pruning) were intersected with state-specific genomic segments by first extracting chromatin-state coordinates with Linux command-line tools (cut and awk) and then identifying SNP–state overlaps using the R package data.table. SNPs falling within each chromatin state were retained. For embryonic stages, SNPs belonging to the same chromatin state across all 11 stages were merged to generate embryonic state-specific SNP sets. For adult muscle, SNPs were grouped according to the muscle-specific chromatin-state segmentation.

For each chromatin state, SNP subsets of varying marker densities were sampled using the --thin-count function in Plink v1.9, with five replicates per density. Predictive performance was evaluated using the same five-round five-fold cross-validation procedure described in Section 2.4.1. Because muscle tissue is expected to be more directly related to growth traits, chromatin-state–filtered SNPs from adult muscle were analyzed separately.

To assess whether improvements were dependent on the statistical model, the chromatin state showing the highest predictive enhancement (state E6 in adult muscle) was additionally evaluated using GBLUP, BayesB, and BayesC, in addition to BayesA.

### 2.5 Cross-population validation of predictive performance

As shown in Section 3.5, SNPs within the muscle bivalent promoter/enhancer (E6) state significantly improved prediction accuracy. To further evaluate their stability and generalizability across populations, the distantly related PLM population (see Table 1 for sample size and trait statistics) was used as an external validation set. The same number of SNPs located in the muscle E6 state were compared with randomly sampled SNPs of equal density.

Before genomic prediction, phenotypes in the reference population were adjusted for population group effects to avoid confounding between population divergence and additive genetic signals. For model training, SNPs from the reference population were partitioned into five subsets for five-fold cross-validation, and the entire procedure was repeated five times, resulting in 25 training iterations in total. In each iteration, four folds were used to estimate marker effects, and the remaining fold served as the internal validation set. After model fitting, the estimated marker effects were directly applied to the genotypes of PLM individuals to compute uncalibrated genomic estimated breeding values (GEBVs), thereby evaluating the transferability of SNP effects without phenotype-based re-estimation in the target population.

Prediction accuracy in the PLM population was calculated as the Pearson correlation between predicted GEBVs and the observed phenotypes, and was further adjusted by dividing by the square root of the trait heritability to account for finite heritability constraints. The same number of E6-state SNPs was compared against randomly sampled SNPs of equal density to evaluate the specificity of functional annotation–guided SNP selection.

### 2.6 Genome-wide association study (GWAS) of SNPs within the muscle bivalent promoter/enhancer (E6) state

To further characterize the functional relevance of SNPs located in the muscle bivalent promoter/enhancer (E6) state and explore their underlying genetic mechanisms, GWAS was conducted using GEMMA v0.98.1 [29]. Group identity was included as a covariate to account for differences among populations. GWAS was performed for body length using a univariate linear mixed model (LMM) to appropriately control for population structure and individual relatedness.

A standardized genomic relationship matrix (GRM) was first computed with the “-gk 2” option in GEMMA according to the following formula:

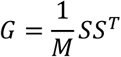

where *M* represents the total number of SNP markers, and S denotes the centered and standardized genotype matrix. The resulting GRM reflects pairwise genetic similarity among individuals and is equivalent to the kinship matrix used in mixed linear model-based GWAS. Subsequently, a univariate LMM was fitted for each SNP using the “-lmm 1” option in GEMMA:

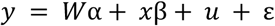

where *y* is the phenotype vector (body length); W is the design matrix for fixed effects, including group identity and the top principal components (PC1 and PC2); α is the corresponding vector of fixed-effect coefficients; x is the genotype vector of the SNP being tested; and β is the SNP effect size. The random polygenic effect is modeled as 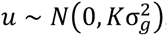, where *K* is the GRM. The residual errors follow 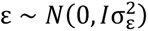. SNP significance was assessed using the Wald test.

Complete GWAS results are provided in **Table S3**. SNP annotation was performed using GenomicFeatures [35] and ChIPseeker [36], followed by Gene Ontology (GO) enrichment analysis using the R package clusterProfiler [40] to identify biological processes and molecular functions enriched among significant SNPs.

To further refine candidate variants and genes associated with growth traits, RNA-seq data from *L. vannamei* muscle were incorporated for functional validation. Raw reads were downloaded from NCBI SRA, with accession numbers SRR6466352, SRR6466355, and SRR6466357 for fast-growth performance (FG) individuals, and SRR6466336, SRR6466339, and SRR6466341 for slow-growth performance (SG) individuals [41]. Quality control and filtering—including automatic adapter removal, trimming of low-quality bases, and minimum-length filtering—were performed using SeqyClean v1.10.09 (https://github.com/ibest/seqyclean), following the preprocessing strategy described in Santos et al. (2021). Reads were further filtered by requiring an average Phred quality ≥ 23, end-base quality ≥ 30, and a minimum read length of 65 bp, consistent with the original workflow. Clean reads were subsequently mapped to the same chromosome-level reference genome used in this study for whole-genome resequencing and CUT&Tag. Read quantification and normalization were performed as in the original study to ensure methodological consistency. Differential expression between the FG and SG groups was assessed with the Wilcoxon rank-sum test (*p* < 0.05), and gene expression visualization was conducted with GraphPad Prism v8.0.2.

## 3. Results

### 3.1 Population characteristics and SNP discovery

The body length and body weight of the 972 *L. vannamei* individuals exhibited normal distributions (**Fig. 1A**), consistent with quantitative traits controlled by multiple genes [42]. Significant differences in both traits were observed among the three populations of the same age (**Fig. 1B**), indicating an effect of genetic background on growth. Linear regression further revealed a strong correlation between body length and body weight (R² = 0.918) **(Fig. S1**).

**Fig. 1.**
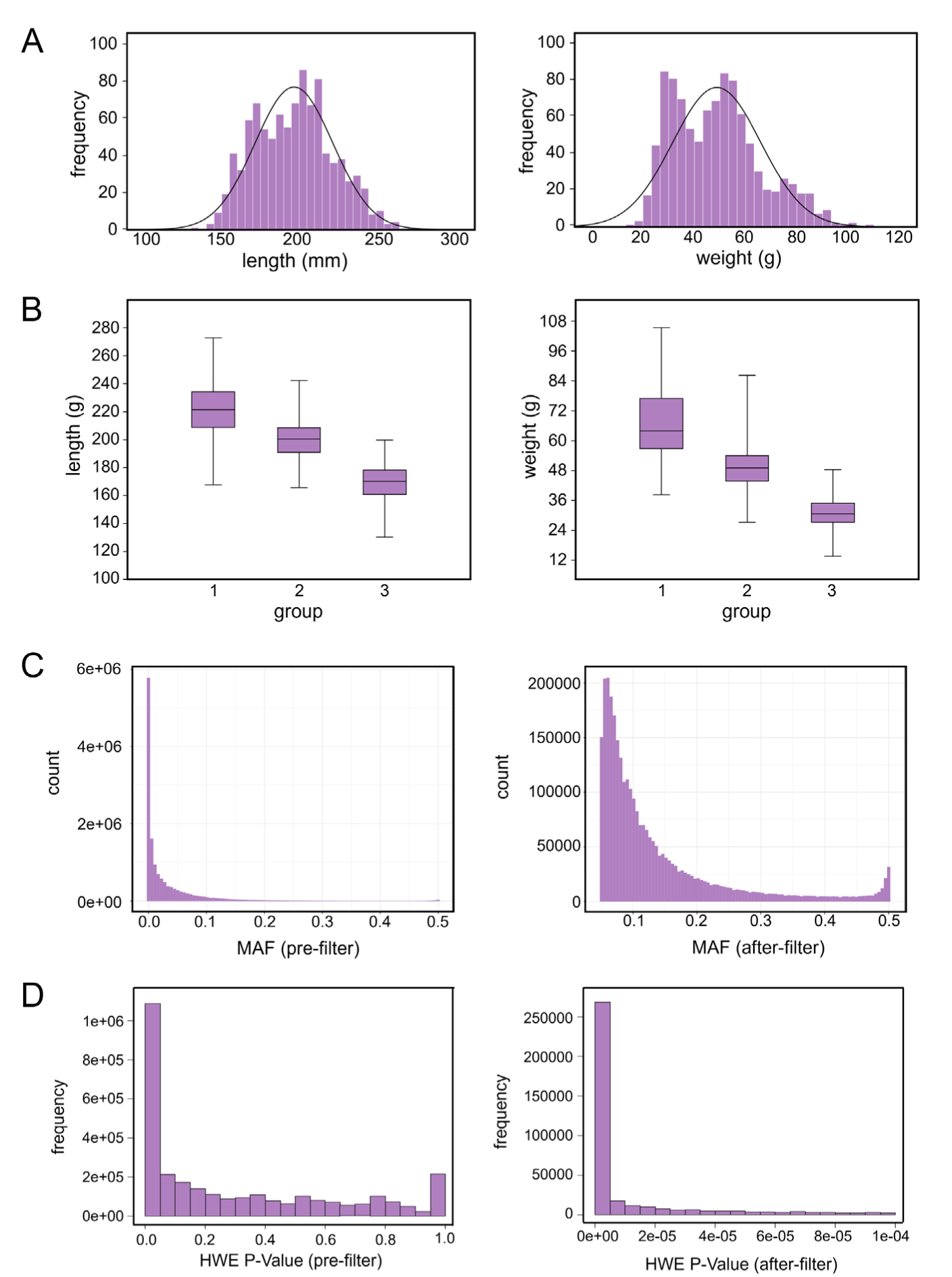
Phenotypic distributions and genotypic quality control for growth traits in selective breeding populations of *Litopenaeus vannamei*. (A) Frequency distributions of body length (left) and body weight (right), both showing normal distributions. (B) Boxplots of body length and body weight for the three populations in the breeding program. (C) Allele frequency distributions before (left) and after (right) filtering for minor allele frequency (MAF) > 0.05. (D) Distributions of Hardy–Weinberg equilibrium (HWE) *p*-values before (left) and after (right) filtering (*p* < 0.0001).

Resequencing of the 972 individuals identified 14,276,173 SNPs after imputation. Following minor allele frequency filtering (MAF < 0.05), 2,978,750 SNPs remained (**Fig. 1C**), and HWE filtering yielded 2,609,549 high-quality SNPs for downstream GS analysis (**Fig. 1D**).

Heritability estimates for body weight and body length were 0.29 ± 0.07 and 0.44 ± 0.08, respectively. Given the higher heritability of body length, it was selected as the representative growth trait for subsequent GS analyses.

### 3.2 BayesA model achieved the highest prediction accuracy for the growth trait in L. vannamei

Principal component analysis (PCA) revealed clear population structure among the samples (**Fig. 2A**). Consistently, ADMIXTURE analysis illustrated distinct ancestry components with varying degrees of admixture (**Fig. S2**), reflecting their divergent breeding histories. The kinship heatmap exhibited a block-like pattern of within-population genetic similarity (**Fig. 2B**). Together, these analyses confirmed the presence of population stratification. To control for this effect while maintaining statistical power, population was included as a fixed effect in subsequent analyses.

**Fig. 2.**
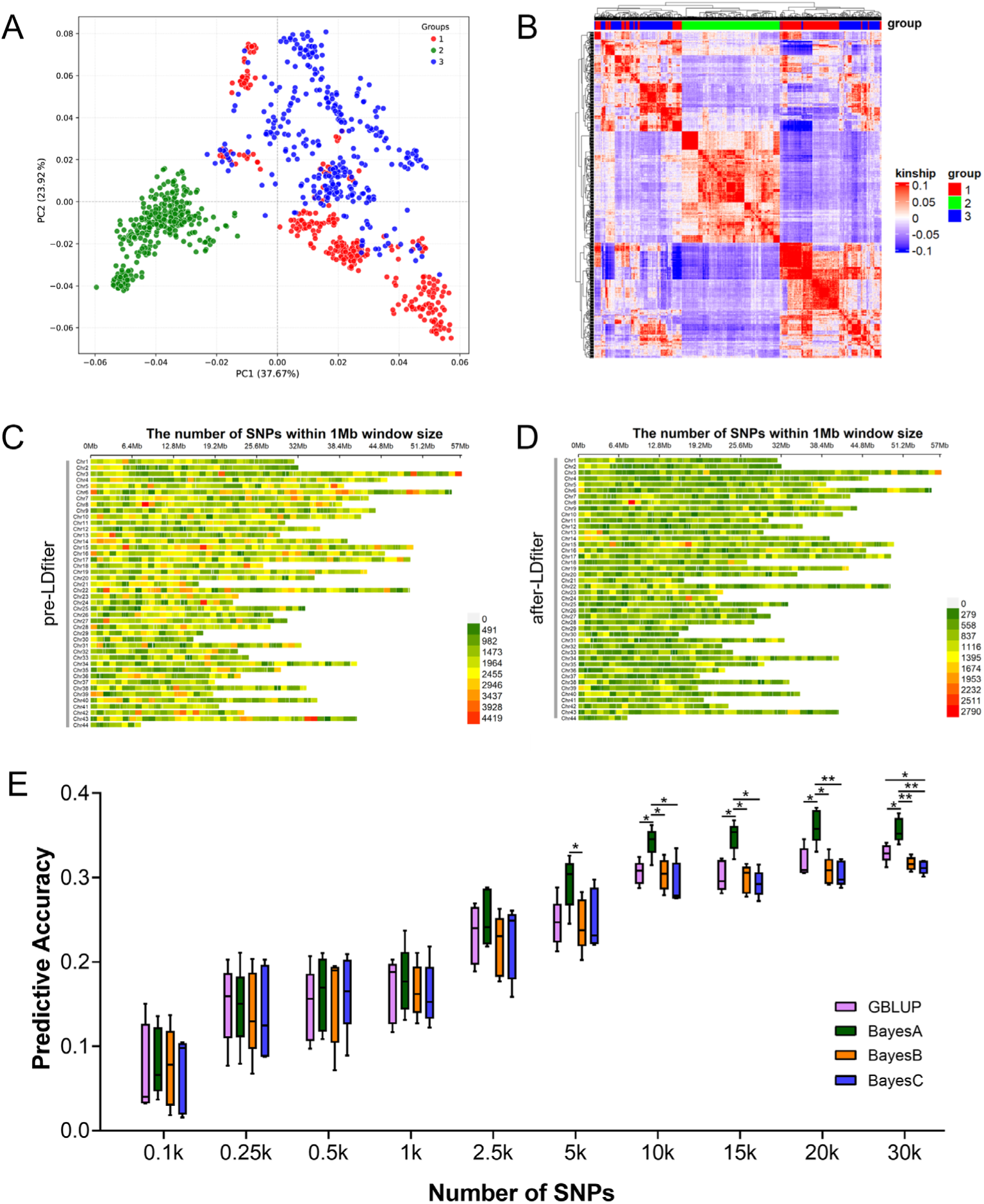
Effects of SNP density and training population size on genomic selection (GS) accuracy for body length in *Litopenaeus vannamei*. (A) Principal component analysis (PCA) based on genome-wide SNPs, showing clear population stratification along PC1 and PC2. (B) Heatmap of the genomic relationship matrix illustrating pairwise relatedness among individuals. (C, D) SNP density plots across chromosomes before and after linkage disequilibrium (LD) pruning, showing even genomic coverage after pruning. (E) Boxplots of predictive accuracy of genomic estimated breeding values (GEBVs) across four GS models (GBLUP, BayesA, BayesB, and BayesC) at different SNP densities. For each density, SNPs were randomly sampled five times and assessed using fivefold cross-validation repeated five times. Accuracy increased with SNP density but plateaued near 10k SNPs, with BayesA consistently achieving the highest accuracy.

After quality control, 2,609,549 SNPs were retained and were evenly distributed across the genome (**Fig. 2C**), consistent with the expectation that most SNPs arise from genetic drift and lack strong regional bias [43]. Following LD filtering, 1,081,117 SNPs remained for GS model training, were also evenly distributed (**Fig. 2D**).

The predictive performance of four GS models (GBLUP, BayesA, BayesB, and BayesC) was compared. Prediction accuracy increased with SNP number across all models, but gains diminished at higher densities, with accuracy plateauing near 10k SNPs (**Fig. 2E; Table S4**). BayesA consistently outperformed the other models under multiple densities (**Fig. 2E; Table S4**) and was therefore adopted for subsequent analyses.

Prediction accuracy and stability also improved with increasing training sample size, reaching a plateau at ∼750 individuals (**Fig. S3; Table S4**). This indicates that the 972 samples used in this study were sufficient to ensure reliable GS analyses and provided a robust foundation for downstream comparisons.

### 3.3 Differential effects of histone modification–based SNP subsets on genomic selection

CUT&Tag data from 11 embryonic stages and adult muscle tissue of *L. vannamei* were analyzed for four histone modifications (H3K4me1, H3K4me3, H3K27ac, H3K27me3) together wih an IgG control to identify SNPs with potential regulatory functions. Sequencing alignment rates exceeded 86% (mostly >90%) (**Table S5**), indicating high data quality. Further quality assessments, including fragment size distribution, peak signal intensity heatmaps, and the genomic feature distribution of histone modification peaks (**Fig. S4**–**S8**), also confirmed that both the newly generated embryonic-stage samples and the adult muscle samples produced reliable CUT&Tag profiles [33,44,45]. The number of histone modification peaks varied substantially across developmental stages and tissues (ranging from 212 to 33,694), reflecting their dynamic regulatory activity, whereas the high concordance between biological replicates demonstrated the robustness of the CUT&Tag signals. The number of peaks detected at each developmental stage or in adult muscle tissue, together with the corresponding numbers of SNPs located within each peak set, are summarized in **Table S6**.

When merging peak regions from all embryonic stages and adult muscle tissue, histone modification signals collectively covered ∼65.34% of annotated coding genes (**Fig. S9**), indicating widespread regulatory element occupancy across the genome had been identified. In total, 306,487 SNPs (11.47% of all high-quality SNPs) were located within histone modification peaks and were broadly distributed across the genome (**Fig. 3A**). Among these, H3K4me3 peaks contained the largest proportion (7.34%), followed by H3K4me1 (4.72%), H3K27me3 (2.15%), and H3K27ac (1.34%) (**Table 2**).

**Fig. 3.**
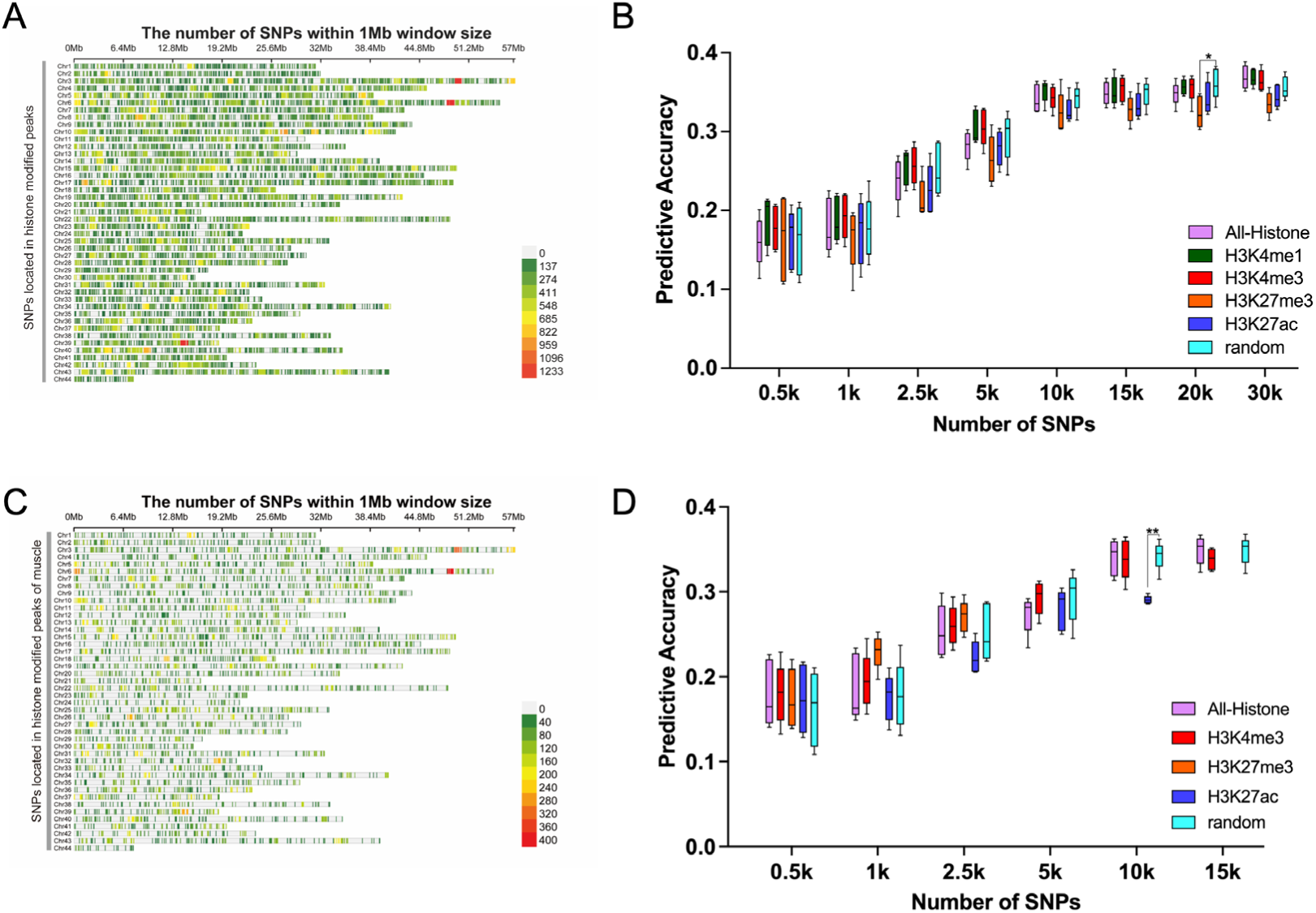
Genomic distribution and predictive performance of histone modification–based SNPs in genomic selection (GS) for growth traits of *Litopenaeus vannamei*. (A) Genomic distribution of SNPs located within histone modification peaks from embryonic stages and adult muscle, showing broad and even coverage. (B) Predictive accuracy of SNP subsets from histone modification peaks of embryonic and muscle tissues at varying marker densities, compared with randomly selected SNPs. For each density, SNPs were sampled five times, and predictive accuracy was assessed with BayesA using five rounds of fivefold cross-validation (25 replicates per density). (C) Genomic distribution of SNPs within histone modification peaks from adult muscle tissue, showing uniform chromosomal coverage. (D) Predictive accuracy of SNP subsets from muscle histone modification peaks compared with random SNPs at the same density. Evaluation procedure were as in (B). Significance levels: \**p* < 0.05, \*\**p* < 0.01.

**Table 2.**
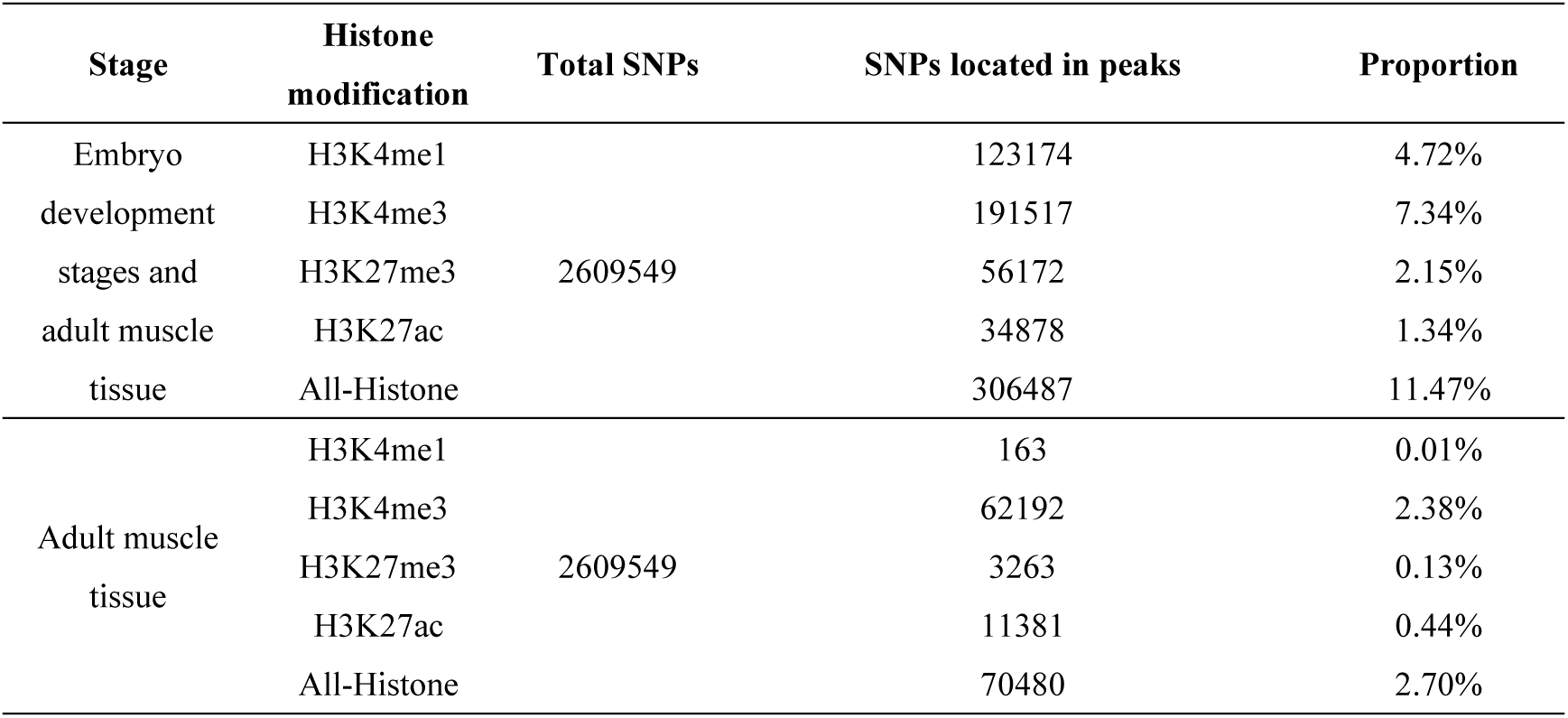
Number and proportion of SNPs located within histone modification peaks.

| Stage | Histone modification | Total SNPs | SNPs located in peaks | Proportion |
| --- | --- | --- | --- | --- |
| Embryo development stages and adult muscle tissue | H3K4me1 | 2609549 | 123174 | 4.72% |
|  | H3K4me3 |  | 191517 | 7.34% |
|  | H3K27me3 |  | 56172 | 2.15% |
|  | H3K27ac |  | 34878 | 1.34% |
|  | All-Histone |  | 306487 | 11.47% |
| Adult muscle tissue | H3K4me1 | 2609549 | 163 | 0.01% |
|  | H3K4me3 |  | 62192 | 2.38% |
|  | H3K27me3 |  | 3263 | 0.13% |
|  | H3K27ac |  | 11381 | 0.44% |
|  | All-Histone |  | 70480 | 2.70% |

To test their utility for GS, SNP subsets of varying densities (0.5k–30k) were sampled from each histone modification region and their union (All-Histone) and compared with random SNPs using BayesA and five-fold cross-validation. Prediction accuracy increased with SNP density and plateaued near 10k. At low densities (0.5k–5k), SNPs from H3K4me1 and H3K4me3 showed higher accuracy (4.00%–18.75%) and greater stability than random SNPs, although differences were not always significant (**Fig. 3B**; **Table 3**). By contrast, SNPs from H3K27me3 and H3K27ac often underperformed, with H3K27me3 accuracy significantly reduced at 20k density. Combining all histone modifications did not improve prediction, likely because broad inclusion diluted tissue- or stage-specific signals. These results suggest that different histone modifications provide distinct functional annotation information, which can differentially influence the prediction accuracy of GS when used for SNP selection.

**Table 3.** Predictive accuracy of SNP subsets from histone modification peaks in embryonic and muscle tissues at varying marker densities.

| Density | Random | H3K4me1 | H3K4m3 | H3K27me3 | H3K27ac | All-Histone |
| --- | --- | --- | --- | --- | --- | --- |
| 0.5k | 0.16 ± 0.04 | 0.19 ± 0.03<br>(+18.75%) | 0.18 ± 0.03<br>(+12.5%) | 0.16 ± 0.05<br>(0%) | 0.16 ± 0.04<br>(0%) | 0.16 ± 0.03<br>(0%) |
| 1k | 0.18 ± 0.04 | 0.19 ± 0.03<br>(+5.56%) | 0.19 ± 0.03<br>(+5.56%) | 0.17 ± 0.04<br>(-5.56%) | 0.17 ± 0.04<br>(-5.56%) | 0.18 ± 0.04<br>(0%) |
| 2.5k | 0.25 ± 0.03 | 0.26 ± 0.02<br>(+4.00%) | 0.26 ± 0.02<br>(+4.00%) | 0.21 ± 0.02<br>(-16.00%) | 0.23 ± 0.03<br>(-8.00%) | 0.24 ± 0.03<br>(-4.00%) |
| 5k | 0.29 ± 0.03 | 0.31 ± 0.02<br>(+6.90%) | 0.31 ± 0.02<br>(+6.90%) | 0.26 ± 0.03<br>(-10.34%) | 0.28 ± 0.02<br>(-3.45%) | 0.28 ± 0.02<br>(-3.45%) |
| 10k | 0.34 ± 0.02 | 0.35 ± 0.02<br>(+2.94%) | 0.34 ± 0.02<br>(0%) | 0.32 ± 0.03<br>(-5.88%) | 0.33 ± 0.02<br>(-2.94%) | 0.34 ± 0.02<br>(0%) |
| 15k | 0.35 ± 0.02 | 0.35 ± 0.02<br>(0%) | 0.35 ± 0.02<br>(0%) | 0.33 ± 0.02<br>(-5.71%) | 0.33 ± 0.02<br>(-5.71%) | 0.35 ± 0.02<br>(0%) |
| 20k | 0.36 ± 0.02 | 0.36 ± 0.01<br>(0%) | 0.36 ± 0.02<br>(0%) | 0.33 ± 0.02*<br>(-8.33%) | 0.34 ± 0.02<br>(-5.56%) | 0.35 ± 0.02<br>(-2.78%) |
| 30k | 0.36 ± 0.01 | 0.37 ± 0.01<br>(+2.78%) | 0.37 ± 0.01<br>(+2.78%) | 0.34 ± 0.01<br>(-5.56%) | 0.35 ± 0.01<br>(-2.78%) | 0.37 ± 0.01<br>(+2.78%) |

Note: Breeding value prediction accuracy is presented as mean ± SD. Percent values in parentheses indicate the relative change in accuracy (increase “+” or decrease “-”) compared with randomly sampled SNPs at the same SNP density. Statistical significance: \* $p < 0.05$ .

Because muscle is directly related to growth, we further examined muscle-specific regulatory regions, identifying 70,480 SNPs (2.70% of all) across the four histone marks (**Fig. 3C**; **Table 2**). The H3K4me1 subset was small (163 SNPs), but at 0.1k density, its prediction accuracy (0.11 ± 0.02) exceeded that of random SNPs (0.08 ± 0.05) by 37.5%, with improved stability (**Table 4**). Across muscle histone marks, accuracy increased with density and plateaued at ∼10k. At low densities (0.5k–2.5k), H3K4me3 subsets consistently outperformed random SNPs (4%–12.5%), while H3K27ac subsets generally underperformed. Notably, H3K27me3 SNPs from muscle showed a trend of improved accuracy (**Fig. 3D**; **Table 4**), in contrast to their performance when embryonic data were included.

**Table 4.** Predictive accuracy of SNP subsets from histone modification peaks in adult muscle tissue at varying marker densities.

| Density | Random | H3K4me1 | H3K4m3 | H3K27me3 | H3K27ac | All-Histone |
| --- | --- | --- | --- | --- | --- | --- |
| 0.1k | 0.08 ± 0.05 | 0.11 ± 0.02<br>(+37.5%) | / | / | / | / |
| 0.5k | 0.16 ± 0.04 | / | 0.18 ± 0.04<br>(+12.5%) | 0.17 ± 0.04<br>(+6.25%) | 0.17 ± 0.04<br>(+6.25%) | 0.18 ± 0.04<br>(+12.5%) |
| 1k | 0.18 ± 0.04 | / | 0.2 ± 0.03<br>(+11.11%) | 0.23 ± 0.02<br>(+27.78%) | 0.18 ± 0.03<br>(0%) | 0.19 ± 0.04<br>(+5.56%) |
| 2.5k | 0.25 ± 0.03 | / | 0.26 ± 0.02<br>(+4.00%) | 0.27 ± 0.02<br>(+8.00%) | 0.22 ± 0.02<br>(-12.00%) | 0.25 ± 0.03<br>(0%) |
| 5k | 0.29 ± 0.03 | / | 0.29 ± 0.02<br>(0%) | / | 0.28 ± 0.02<br>(-3.45%) | 0.27 ± 0.02<br>(-6.90%) |
| 10k | 0.34 ± 0.02 | / | 0.34 ± 0.02<br>(0%) | / | 0.29 ± 0.01**<br>(-14.71%) | 0.34 ± 0.02<br>(0%) |
| 15k | 0.35 ± 0.02 | / | 0.34 ± 0.01<br>(-2.86%) | / | / | 0.35 ± 0.02<br>(0%) |

Note: Breeding value prediction accuracy is presented as mean ± SD. Percent values in parentheses indicate the relative change in accuracy (increase “+” or decrease “-”) compared with randomly sampled SNPs at the same SNP density. Statistical significance: \*\* $p < 0.01$ .

Overall, histone modification–based SNP selection improved the predictive accuracy and stability of GS models under low marker densities, though effects varied by tissue and modification. H3K4me3 consistently provided stable gains, supporting its role as a key regulatory mark. Using all tissues together yielded modest improvements, whereas muscle-specific information produced stronger trait-related benefits. These findings suggest that integrating histone modification data to refine functional SNP annotations may further enhance GS accuracy.

### 3.4 Functional annotation of embryonic chromatin states reveals SNP subsets with distinct predictive power

The GS results based on individual histone modifications indicated that SNPs located in different functional regions exert distinct predictive effects. To obtain higher-resolution functional information, we integrated four histone modification datasets (H3K4me1, H3K4me3, H3K27me3, and H3K27ac) from embryonic stages and adult muscle of *L. vannamei* and identified eight chromatin states using ChromHMM [17]. These included: flanking bivalent transcription start site (TSS)/enhancer 1 (BivFInk1), quiescent/low expression regions (Quies), weak repressed Polycomb regions (ReprPCWk), repressed Polycomb regions (ReprPC), weak enhancer regions (EnhWk), flanking bivalent TSS/enhancer 2 (BivFInk2), weak transcription regions (TxWk), and flanking TSS downstream regions (TssFlnkD) (**Fig. 4A**). As histone modification effects appeared tissue-specific, chromatin states from embryonic stages and adult muscle were analyzed separately.

**Fig. 4.**
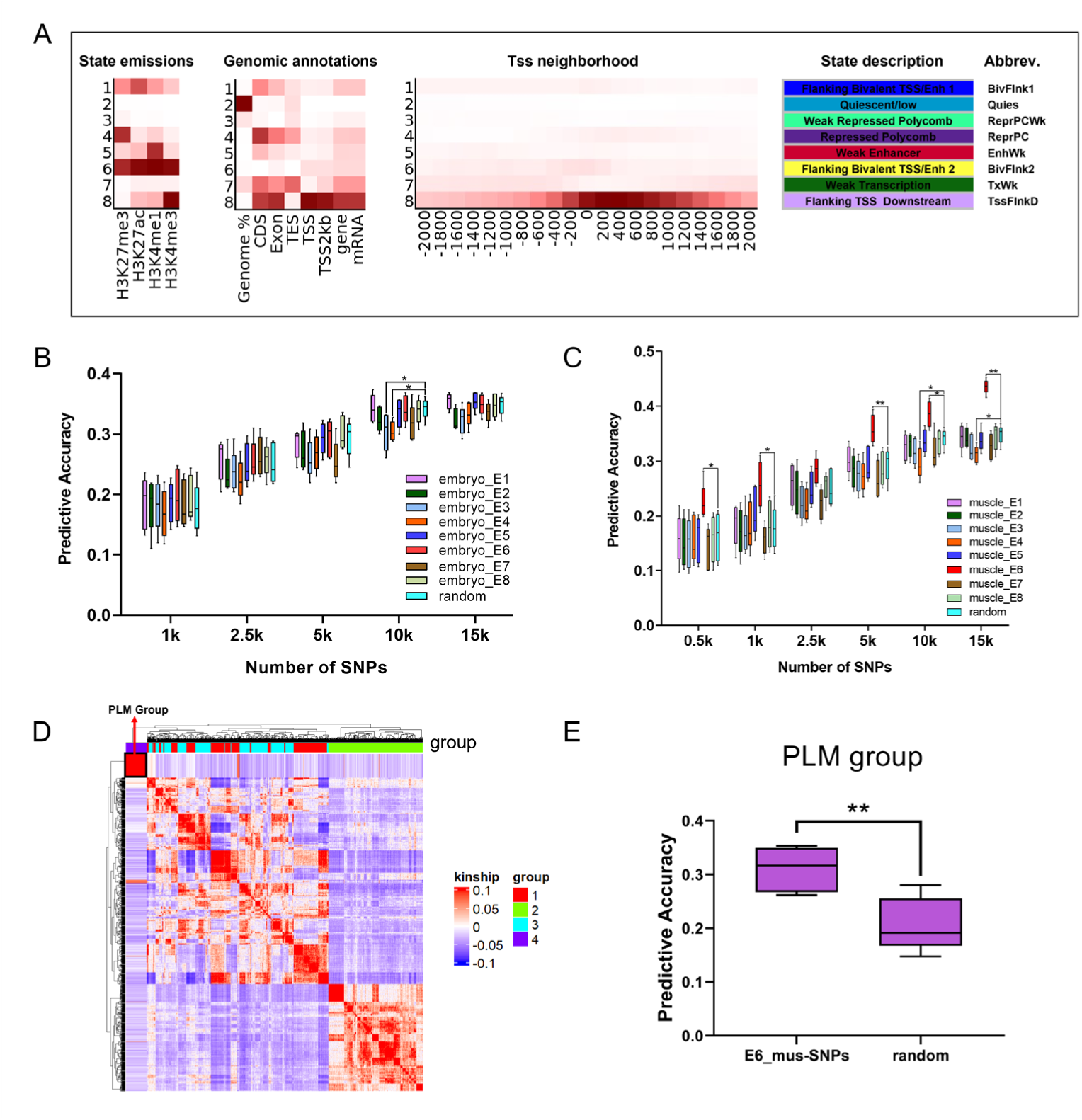
Chromatin state–based SNP annotation reveals tissue-specific predictive performance and cross-population robustness in genomic selection (GS) for *Litopenaeus vannamei* growth traits. (A) Chromatin state annotation based on H3K4me1, H3K4me3, H3K27me3, and H3K27ac marks from the embryonic and muscle tissues. The left panel shows the emission parameters of the ChromHMM 8-state model, where color intensity represents the probability of each histone modification occurring within a given state. The middle panel depicts state enrichments across external genomic annotations, with color gradients indicating enrichment levels. The right panel illustrates fold enrichment of chromatin states within ± 2 kb of transcription start sites (TSSs) at 200 bp resolution, with deeper red reflecting stronger enrichment. The state descriptions and abbreviations are summarized in the right panel. (B) Predictive accuracy of SNPs located in embryonic chromatin states across varying marker densities compared with randomly sampled SNPs. (C) Predictive accuracy of SNPs located in muscle chromatin states. SNPs within the muscle bivalent promoter/enhancer state (E6) consistently outperformed random SNPs across densities, achieving significantly higher predictive accuracy. (D) Kinship matrix showing genetic divergence between the training population and the independent PLM validation population. (E) Cross-population validation of muscle E6 SNPs in the PLM population. Predictive accuracy was compared with equal-sized sets of random SNPs. Models were trained with fivefold cross-validation in the reference population, with PLM as the validation set. E6 SNPs improved accuracy by 47.62% (from 0.21 ± 0.05 to 0.31 ± 0.04). Significance levels: \**p* < 0.05, \*\**p* < 0.01.

In embryos, the number of chromatin state sites ranged from 11,313 to 113,769, with SNP coverage from 13,291 to 1,912,007, reflecting dynamic changes in chromatin accessibility and its relationship with genetic variation (**Table S7**). The quiescent/low expression state (Quies, E2), which lacks histone modification marks and likely represents inactive regions, encompassed the largest number of SNPs (90.10%). In contrast, SNPs were far fewer in active states such as TxWk (E7), BivFInk1/2 (E1/E6), and TssFlnkD (E8) (**Table 5**).

**Table 5.** Number and proportion of SNPs located within different chromatin state regions in embryonic tissues.

| Chromatin state | Total SNPs | SNPs in chromatin state regions | Proportion |
| --- | --- | --- | --- |
| E1 BivFlnk1 | 2609549 | 201729 | 7.73% |
| E2 Quies |  | 2351162 | 90.10% |
| E3 ReprPCWk |  | 828478 | 31.75% |
| E4 ReprPC |  | 238452 | 9.14% |
| E5 EnhWk |  | 277111 | 10.62% |
| E6 BivFlnk2 |  | 107027 | 4.10% |
| E7 TxWk |  | 1080973 | 41.42% |
| E8 TssFlnkD |  | 312303 | 11.97% |

To evaluate predictive performance, SNP subsets (0.5k–30k, five replicates per density) were sampled from each chromatin state and assessed with the BayesA model using five rounds of five-fold cross-validation. SNPs in BivFlnk1 (E1), EnhWk (E5), BivFlnk2 (E6), and TssFlnkD (E8) improved prediction accuracy by 4.00–11.11% over random SNPs at low-to-moderate densities (1k– 2.5k) (**Fig. 4B**; **Table 6**). However, the overall improvement from embryonic chromatin states was modest, likely because embryonic regulatory landscapes differ from those driving adult growth phenotypes.

**Table 6.**
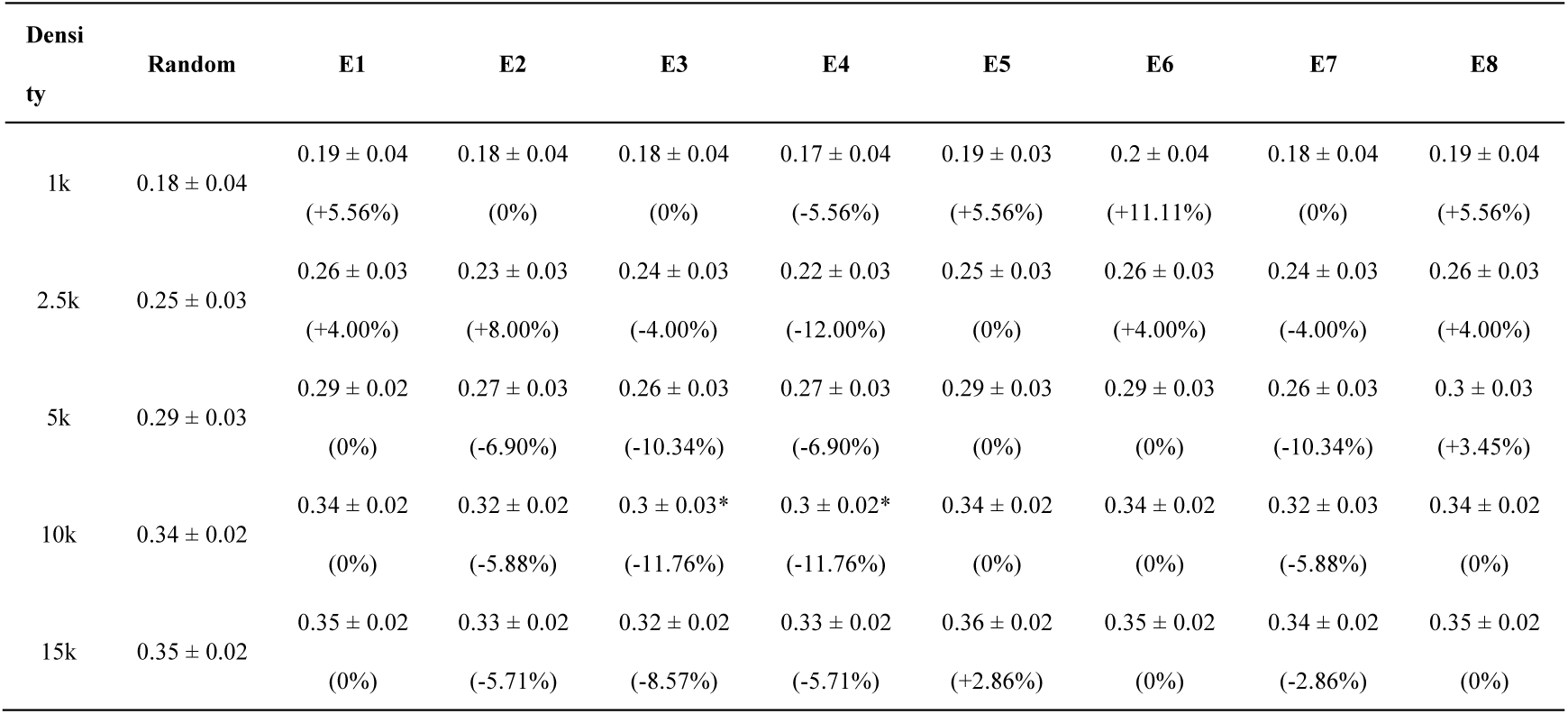
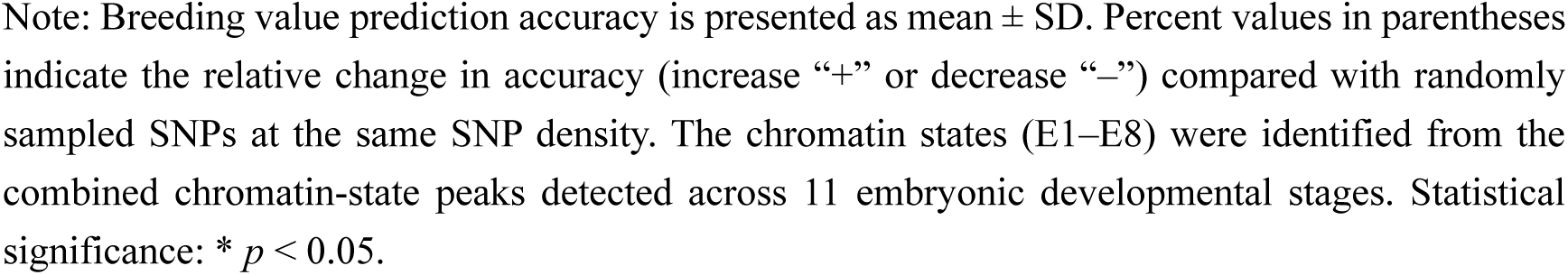
Predictive accuracy of SNPs located in embryonic chromatin states across varying marker densities compared with randomly sampled SNPs.

| Density | Random | E1 | E2 | E3 | E4 | E5 | E6 | E7 | E8 |
| --- | --- | --- | --- | --- | --- | --- | --- | --- | --- |
| 1k | 0.18 ± 0.04 | 0.19 ± 0.04<br>(+5.56%) | 0.18 ± 0.04<br>(0%) | 0.18 ± 0.04<br>(0%) | 0.17 ± 0.04<br>(-5.56%) | 0.19 ± 0.03<br>(+5.56%) | 0.2 ± 0.04<br>(+11.11%) | 0.18 ± 0.04<br>(0%) | 0.19 ± 0.04<br>(+5.56%) |
| 2.5k | 0.25 ± 0.03 | 0.26 ± 0.03<br>(+4.00%) | 0.23 ± 0.03<br>(+8.00%) | 0.24 ± 0.03<br>(-4.00%) | 0.22 ± 0.03<br>(-12.00%) | 0.25 ± 0.03<br>(0%) | 0.26 ± 0.03<br>(+4.00%) | 0.24 ± 0.03<br>(-4.00%) | 0.26 ± 0.03<br>(+4.00%) |
| 5k | 0.29 ± 0.03 | 0.29 ± 0.02<br>(0%) | 0.27 ± 0.03<br>(-6.90%) | 0.26 ± 0.03<br>(-10.34%) | 0.27 ± 0.03<br>(-6.90%) | 0.29 ± 0.03<br>(0%) | 0.29 ± 0.03<br>(0%) | 0.26 ± 0.03<br>(-10.34%) | 0.3 ± 0.03<br>(+3.45%) |
| 10k | 0.34 ± 0.02 | 0.34 ± 0.02<br>(0%) | 0.32 ± 0.02<br>(-5.88%) | 0.3 ± 0.03*<br>(-11.76%) | 0.3 ± 0.02*<br>(-11.76%) | 0.34 ± 0.02<br>(0%) | 0.34 ± 0.02<br>(0%) | 0.32 ± 0.03<br>(-5.88%) | 0.34 ± 0.02<br>(0%) |
| 15k | 0.35 ± 0.02 | 0.35 ± 0.02<br>(0%) | 0.33 ± 0.02<br>(-5.71%) | 0.32 ± 0.02<br>(-8.57%) | 0.33 ± 0.02<br>(-5.71%) | 0.36 ± 0.02<br>(+2.86%) | 0.35 ± 0.02<br>(0%) | 0.34 ± 0.02<br>(-2.86%) | 0.35 ± 0.02<br>(0%) |

Note: Breeding value prediction accuracy is presented as mean ± SD. Percent values in parentheses indicate the relative change in accuracy (increase “+” or decrease “-”) compared with randomly sampled SNPs at the same SNP density. The chromatin states (E1–E8) were identified from the combined chromatin-state peaks detected across 11 embryonic developmental stages. Statistical significance: \* $p < 0.05$ .

**Table 7.** Number and proportion of SNPs located within different chromatin state regions of adult shrimp muscle tissue.

| Chromatin state | Total SNPs | SNPs in chromatin state regions | Proportion |
| --- | --- | --- | --- |
| E1 BivFlnk1 | 2609549 | 44299 | 1.70% |
| E2 Quies |  | 1689212 | 64.73% |
| E3 ReprPCWk |  | 207586 | 7.95% |
| E4 ReprPC |  | 39606 | 1.52% |
| E5 EnhWk |  | 17290 | 0.66% |
| E6 BivFlnk2 |  | 15073 | 0.58% |
| E7 TxWk |  | 467392 | 17.91% |
| E8 TssFlnkD |  | 129889 | 4.98% |

Notably, SNPs from repressive or weak transcriptional states reduced predictive performance. In ReprPCWk (E3), ReprPC (E4), and TxWk (E7), accuracy decreased by 2.86–12.00% compared with random SNPs at 2.5k–15k densities, suggesting that such regions contribute little—or even introduce noise—to GS predictions (**Fig. 4B**; **Table 6**).

### Predictive superiority and cross-population robustness of muscle chromatin state–based SNPs

Analysis of chromatin states in adult shrimp muscle revealed 19,763–105,594 peaks across different states, reflecting substantial variation among functional regions such as promoters, enhancers, and repressive domains. Corresponding SNP coverage ranged from 15,073 to 1,689,212, with the quiescent state (E2) again encompassing the largest number of SNPs (**Tables 7, S7**).

Genomic prediction analyses revealed that SNPs within the muscle bivalent promoter/enhancer state (E6) consistently exhibited the strongest predictive performance across all marker densities. At low densities (0.5k–2.5k), E6 SNPs improved prediction accuracy by 12.00–44.44% compared with random LD-filtered SNPs. At a density of 5k, E6 SNPs achieved accuracy comparable to that obtained using 15k randomly sampled SNPs. At 15k density, E6 SNPs produced an additional 25.71% improvement in accuracy relative to random SNPs (**Fig. 4C**; **Table 8**).

**Table 8.** Predictive accuracy of SNPs located in adult muscle chromatin states across varying marker densities compared with randomly sampled SNPs.

| Density | Random | E1 | E2 | E3 | E4 | E5 | E6 | E7 | E8 |
| --- | --- | --- | --- | --- | --- | --- | --- | --- | --- |
| 0.5k | 0.16 ± 0.04 | 0.16 ± 0.04<br>(0%) | 0.15 ± 0.04<br>(-6.25%) | 0.15 ± 0.05<br>(-6.25%) | 0.16 ± 0.04<br>(0%) | 0.16 ± 0.04<br>(0%) | 0.23 ± 0.03*<br>(+43.75%) | 0.14 ± 0.04<br>(-12.5%) | 0.16 ± 0.04<br>(0%) |
| 1k | 0.18 ± 0.04 | 0.19 ± 0.04<br>(+5.56%) | 0.17 ± 0.04<br>(-5.56%) | 0.17 ± 0.04<br>(-5.56%) | 0.18 ± 0.04<br>(0%) | 0.21 ± 0.04<br>(+16.67%) | 0.26 ± 0.04*<br>(+44.44%) | 0.16 ± 0.03<br>(-11.11%) | 0.18 ± 0.04<br>(0%) |
| 2.5k | 0.25 ± 0.03 | 0.26 ± 0.04<br>(+3.85%) | 0.24 ± 0.04<br>(-4.00%) | 0.22 ± 0.03<br>(-12.00%) | 0.22 ± 0.03<br>(-12.00%) | 0.25 ± 0.03<br>(0%) | 0.28 ± 0.03<br>(+12.00%) | 0.22 ± 0.03<br>(-12.00%) | 0.26 ± 0.03<br>(+4.00%) |
| 5k | 0.29 ± 0.03 | 0.3 ± 0.03<br>(+3.45%) | 0.28 ± 0.03<br>(-3.45%) | 0.27 ± 0.03<br>(-6.90%) | 0.27 ± 0.03<br>(-6.90%) | 0.29 ± 0.03<br>(0%) | 0.36 ± 0.03**<br>(+24.14%) | 0.27 ± 0.03<br>(-6.90%) | 0.29 ± 0.03<br>(0%) |
| 10k | 0.34 ± 0.02 | 0.33 ± 0.02 (-<br>2.94%) | 0.33 ± 0.02<br>(-2.94%) | 0.32 ± 0.02<br>(-5.88%) | 0.3 ± 0.03*<br>(-11.76%) | 0.34 ± 0.02<br>(0%) | 0.39 ± 0.02*<br>(+14.71%) | 0.32 ± 0.03<br>(-5.88%) | 0.34 ± 0.02<br>(0%) |
| 15k | 0.35 ± 0.02 | 0.35 ± 0.02<br>(0%) | 0.34 ± 0.02<br>(-2.86%) | 0.32 ± 0.02<br>(-8.57%) | 0.31 ± 0.02*<br>(-11.43%) | 0.34 ± 0.02<br>(-2.86%) | 0.44 ± 0.01**<br>(+25.71%) | 0.33 ± 0.02<br>(-5.71%) | 0.35 ± 0.02<br>(0%) |

Note: Breeding value prediction accuracy is presented as mean ± SD. Percent values in parentheses indicate the relative change in accuracy (increase “+” or decrease “-”) compared with randomly sampled SNPs at the same SNP density. The chromatin states (E1–E8) were identified from the combined chromatin-state peaks detected across muscle tissue of adult shrimp. Statistical significance: \* $p < 0.05$ , \*\* $p < 0.01$ .

To further assess the robustness of the chromatin state–based improvement, three additional genomic prediction models (BayesB, BayesC, and GBLUP) were applied to the muscle E6 SNP set. Consistent with the BayesA results, E6 SNPs yielded higher predictive accuracy across all marker densities (Fig. S10), indicating that the improvement in prediction accuracy was not dependent on the prior assumptions of the BayesA model but rather stemmed from the unique biological relevance of the E6 chromatin state. These results demonstrate that E6 SNPs offer strong predictive power and meaningful biological interpretability under both low- and moderate-density scenarios.

In cross-population validation, using the PLM population as an independent validation set, the prediction accuracy of E6 SNPs was significantly improved by 47.62% compared with random SNPs (from 0.21 ± 0.05 to 0.31 ± 0.04, *p* < 0.05) (**Figs. 4D and 4E**), highlighting their excellent generalizability and practical value across populations.

Notably, SNPs within certain chromatin state regions exerted a negative impact on predictive performance. For example, in the quiescent state (E2), weak/strong repressive regions (ReprPCWk, E3; ReprPC, E4), and weak transcriptional activity regions (TxWk, E7), prediction accuracy decreased by 2.86%–12.00% compared with random SNPs under 2.5k–15k marker densities (**Fig. 4C**; **Table 8**), suggesting that regions with low activity, repressive function, or weak transcriptional activity may contribute little, or even introduce noise, to GS prediction.

In summary, SNPs from the E6 (bivalent promoter/enhancer) region exhibited not only high predictive performance within the focal population but also strong robustness in cross-population validation, underscoring their potential as functional genetic markers for application in molecular breeding.

### 3.5 Muscle bivalent promoter/enhancer (E6) state SNPs associated genes are significantly enriched in growth-related pathways

Given the significant advantage of SNPs within the muscle bivalent promoter/enhancer (E6) state in genomic prediction, we further evaluated their direct association with body length to identify potential key regulatory elements, functional genes, and related pathways. The results showed that the heritability of this trait was 0.40, indicating a moderately high level and strong potential for genetic improvement.

A total of 15,073 high-quality SNPs located within the E6 state were included in the GWAS, and these SNPs were relatively evenly distributed across chromosomes (**Fig. 5A**). In total, 751 SNPs showed significant associations with body length (**Fig. 5B**). GO enrichment analysis indicated that the significant SNPs were predominantly enriched in biological processes related to muscle cell differentiation and development, chromatin modification, biosynthesis, gene expression regulation, and metabolic processes (**Fig. 5C**). Enrichment of muscle development pathways underscores their central roles in muscle growth and functional maintenance [46]. The enrichment of chromatin modification pathways revealed the bridging role of epigenetic mechanisms between genetic variation and trait expression [47]. Meanwhile, the enrichment of metabolism-related pathways suggests that these SNPs may indirectly regulate growth performance by affecting cellular metabolism and energy supply [48].

**Fig. 5.**
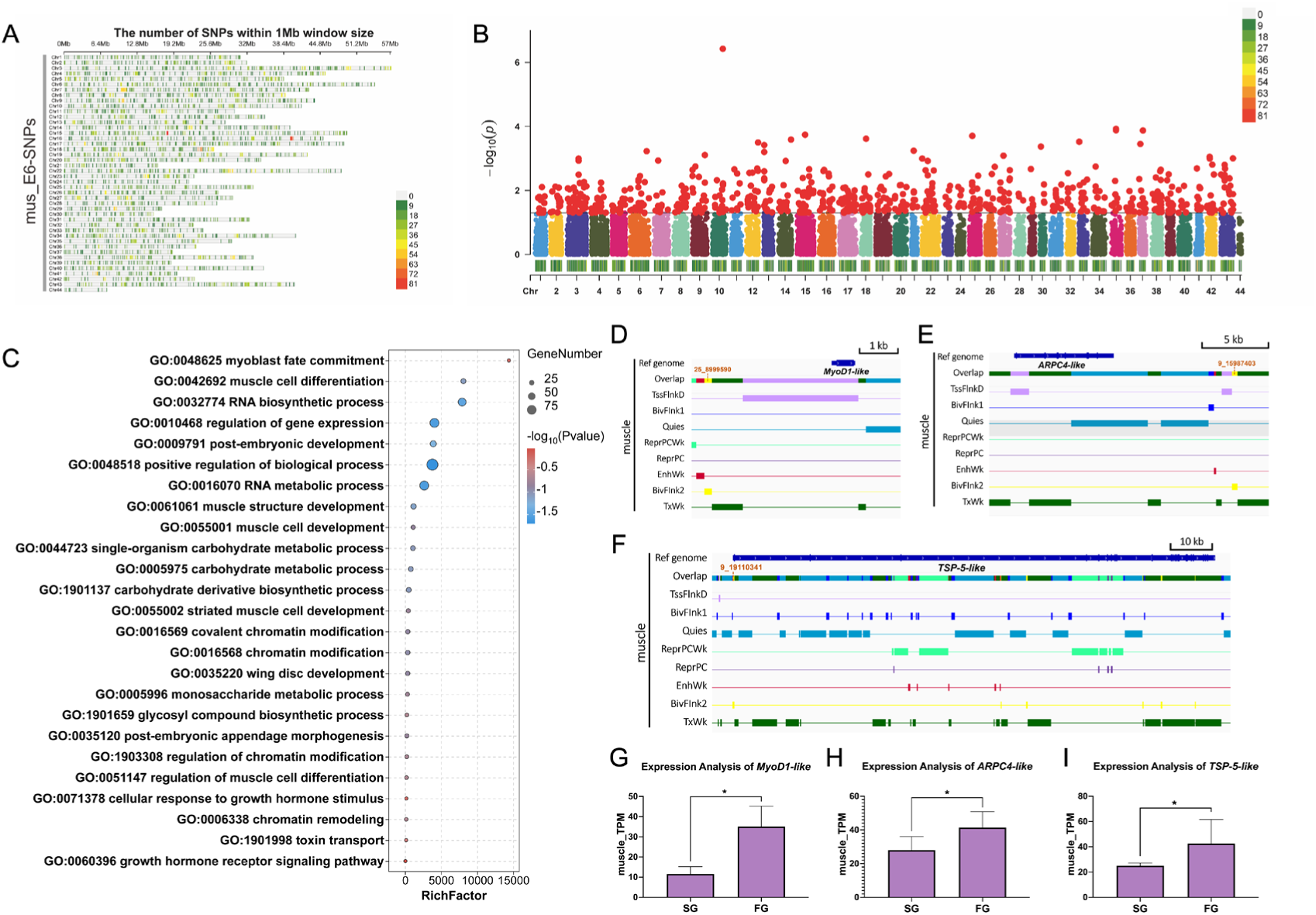
Genome-wide association analysis (GWAS) of SNPs located in the muscle bivalent promoter/enhancer (E6) state. (A) Genomic distribution of 15,073 high-quality E6 SNPs across chromosomes, showing broad and even coverage. (B) Manhattan plot of GWAS for body length, with the horizontal line indicating the significance threshold (*p* < 0.05). Red dots represent significant SNPs, and chromosome density is shown at the bottom. (C) GO functional enrichment of genes annotated from significant E6 SNPs. Bubble size represents the number of enriched genes, and color intensity reflects statistical significance (*p* < 0.05). Results indicate enrichment in multiple growth-related pathways. (D–F) The ChromHMM IGV visualization results of loci 25_8999590, 22_15987403, and 9_19110341, all of which are located within the E6 state region. (G–I) Expression levels of *MyoD1-like*, *ARPC4-like*, and *TSP-5-like* in muscle tissues of fast-growth (FG) and slow-growth (SG) groups, with significantly higher expression in FG shrimp (*p* < 0.05).

Integrating GWAS signals with gene annotation further identified several strong candidate genes associated with growth and their putative regulatory SNPs. For example, XM_027356510.1 (*MyoD1-like*) encodes a master transcription factor essential for myogenic determination [49]. XM_027359869.1 (*ARPC4-like*) encodes a core subunit of the Arp2/3 complex, an actin polymerization factor involved in muscle fiber formation [50]. XM_027375747.1 (*COMP/TSP-5-like*) belongs to the thrombospondin (TSP) family and regulates the development of the muscle– tendon junction [51]. Significant SNPs adjacent to these genes showed strong associations with body length in GWAS (**Fig. 5D–F**), suggesting their important roles in growth regulation. Moreover, transcriptome analysis confirmed that these genes were widely expressed in muscle tissue, with significantly higher expression in the fast-growth (FG) group compared to the slow-growth (SG) group (**Fig. 5G–I**), further supporting their functional roles in growth regulation.

In summary, functional SNPs within the E6 state not only performed well in genomic selection prediction but also revealed a series of growth-related genes and pathways through GWAS, confirming their potential value as functional markers for molecular breeding.

## 4. Discussion

The use of functional genomic annotation to enhance genomic selection (GS) accuracy has long been considered a promising direction [10,14,19,52]. However, practical applications remain scarce in aquaculture species, largely due to the lack of systematic and high-resolution functional annotations. This study provides one of the first systematic demonstrations of integrating histone modification–based functional annotations into genomic selection for shrimp growth traits. By leveraging CUT&Tag-derived high-resolution maps of four histone marks (H3K4me1, H3K4me3, H3K27me3, and H3K27ac), we identified and annotated regulatory elements, and used them to guide SNP selection for GS of growth traits. Our findings demonstrate that epigenetic annotation– based SNP subsets improved prediction accuracy and stability, with statistically significant gains in multiple conditions, particularly SNPs located in the muscle-specific bivalent promoter/enhancer (E6) state. These SNPs not only achieved comparable or superior predictive accuracy under reduced marker densities, but also exhibited strong cross-population generalizability. This strategy provides a feasible path to lowering genotyping costs while maintaining or enhancing GS performance.

### Genetic architecture and the need for functional SNP prioritization

The heritability estimates for body length (0.44) and body weight (0.29) indicated moderate genetic control, consistent with previous reports in aquatic species [53–55]. Among the GS models tested, BayesA achieved the highest accuracy, corroborating earlier findings that it better captures polygenic architectures by allowing heterogeneous effect distributions [2,56].

Traditional GS relies on dense genome-wide SNP coverage, but the inclusion of numerous non-informative markers dilutes the effects of causal variants and rapidly diminishes the marginal benefits of increasing SNP density [3,57,58]. In this study, prediction accuracy saturated at ∼10k SNPs, highlighting the potential of low-density panels optimized by functional annotation for cost-effective breeding applications.

Our results confirmed that SNPs within active regulatory regions such as promoters (H3K4me3, H3K27ac) and enhancers (H3K4me1) conferred moderate accuracy improvements at low densities, consistent with their role in transcriptional regulation. However, the gains from single histone marks were limited, as each modification reflects only a single dimension of chromatin regulation [47,59,60]. In contrast, ChromHMM-based chromatin state annotation integrates multiple marks to partition the genome into biologically meaningful states, thereby improving regulatory element identification and predictive utility.

### Tissue-specific chromatin states and GS performance

GS accuracy is strongly influenced by the genetic relatedness between training and validation populations and often declines in cross-population scenarios [3,58]. Identifying biologically stable, functionally relevant SNPs is therefore essential. Previous studies have attempted to incorporate functional annotations (e.g., genomic structural features) or to up-weight SNPs identified as significant in GWAS, but these approaches generally rely on simplified functional partitions or limited prior knowledge, which restricts their ability to comprehensively capture key regulatory variants [6,61,62]. Our analyses revealed that embryonic chromatin state–based SNPs provided only modest improvements, likely due to developmental-stage differences from adult phenotypes. In contrast, muscle chromatin state SNPs, particularly those in the E6 bivalent promoter/enhancer state, exhibited remarkable predictive performance. At low-to-moderate densities, E6 SNPs achieved accuracy comparable to or exceeding that of 15k random SNPs, and in the independent PLM population, they improved prediction accuracy by 47.62%. This highlights the dual value of E6 SNPs in enhancing both cost-efficiency and cross-population robustness. These results emphasize the importance of tissue-specific regulatory information for identifying functional SNPs that enhance both cost-efficiency and cross-population robustness.

### Biological relevance of E6 SNPs

The superior performance of E6-state SNPs reflects both the intrinsic regulatory properties of bivalent chromatin and the direct functional relevance of variants residing in these regions. Bivalent promoter/enhancer elements marked simultaneously by H3K4me3 and H3K27me3 are well known to maintain key developmental genes in a poised but responsive state, enabling rapid transcriptional activation under environmental or physiological cues [63,64]. This chromatin flexibility facilitates dynamic responses to stressors such as temperature fluctuations, hypoxia, and pathogen exposure— conditions frequently encountered in aquaculture and known to influence growth performance [65– 67]. Consistent with this regulatory potential, our GWAS of 15,073 E6-state SNPs identified 751 significant variants associated with body length, and the associated genes were enriched in pathways related to muscle development, chromatin regulation, transcriptional control, and metabolic processes—core biological functions underlying growth (**Fig. 5C**) (Braun and Gautel, 2011; Zhang et al., 2020; Stitt et al., 2010). Moreover, major candidates such as MyoD1-like, ARPC4-like, and TSP-5-like not only carried significant E6 variants but also showed higher expression in fast-growing individuals (**Fig. 5G**–**I**), providing strong evidence that E6-state SNPs capture regulatory polymorphisms influencing core growth genes.

Importantly, the predictive advantage of E6-state SNPs was model-independent. Across four genomic prediction models—BayesA, BayesB, BayesC, and GBLUP—E6 SNPs consistently outperformed randomly selected SNPs across all marker densities, indicating that their improvement does not rely on the statistical assumptions of any single model. Instead, the enhanced accuracy stems from the underlying biological informativeness of E6-state regulatory elements, highlighting the robustness of chromatin-state–guided SNP prioritization across analytical frameworks and its practical utility for breeding applications.

Building upon this mechanistic and empirical foundation, we propose an epigenome-guided SNP prioritization framework (**Fig. 6**), which integrates multiple histone modification profiles to define key regulatory chromatin states and systematically enrich SNPs within active promoters, enhancers, and bivalent regions. By focusing on genomic segments with high regulatory potential, this framework ensures effective coverage of core functional variants while substantially reducing marker density requirements. Moreover, because regulatory architectures of essential developmental and metabolic pathways tend to exhibit higher evolutionary and population-level stability, SNPs within these regions also retain predictive power across genomes with diverged LD structures. This is consistent with the substantial 47.62% improvement observed in the genetically distant PLM validation population, demonstrating the strong cross-population robustness of E6-state SNPs.

**Fig. 6.**
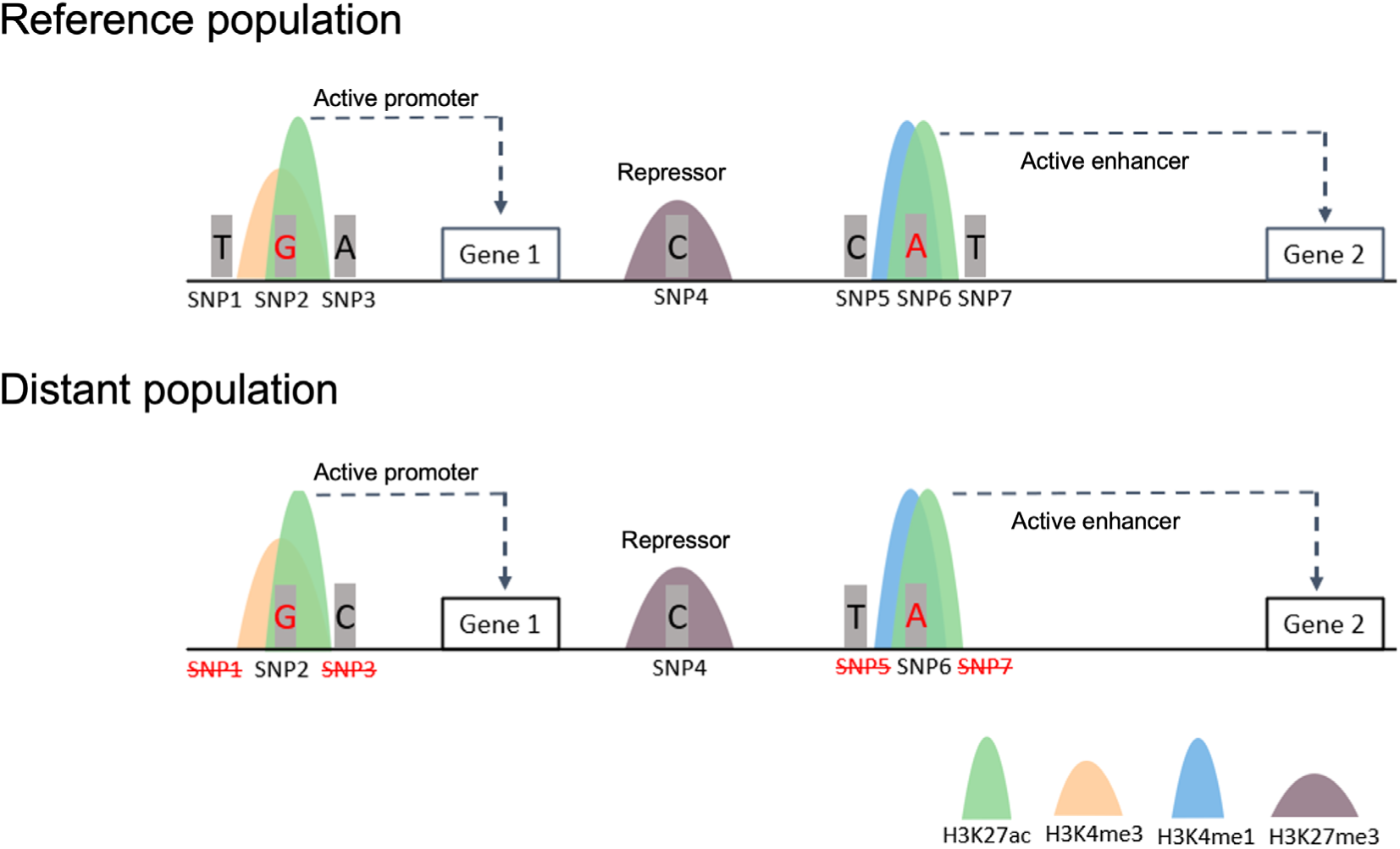
Conceptual framework of histone modification–guided SNP prioritization for improving genomic prediction and cross-population robustness. By integrating chromatin states defined by histone modifications (H3K27ac, H3K4me3, H3K4me1, and H3K27me3), active promoters, enhancers, repressive regions, and bivalent elements can be precisely annotated. SNPs located within these regulatory regions are prioritized as functionally relevant markers, ensuring effective coverage of key regulatory elements. This strategy not only improves the accuracy of genomic selection within populations but also enhances prediction stability and robustness across genetically distant populations, as a higher proportion of phenotype-associated causal SNPs are retained.

Overall, these results demonstrate that chromatin-state–based functional annotation—especially leveraging bivalent promoter/enhancer regions in the adult muscle tissue for the growth trait— provides a biologically grounded and cross-population–robust strategy for enhancing genomic prediction in aquaculture breeding.

### Broader implications for livestock genomic prediction

Although validated here in shrimp, our findings support a broader principle that epigenome-informed regulatory SNP prioritization can enhance genomic prediction across farmed animal species. In livestock species like pigs and cattle, large-scale functional annotation projects (e.g., FAANG) have revealed that causal variants for complex traits are disproportionately enriched in regulatory regions rather than coding sequences [11,18]. Our results extend this understanding by highlighting the specific utility of bivalent domains (marked by H3K4me3 and H3K27me3). Biologically, these domains maintain developmental genes in a "poised" state, allowing for rapid activation in response to environmental stimuli [14,63], a mechanism that is evolutionarily conserved across metazoans [68,69].

In livestock production, animals frequently face variable stressors such as heat stress in dairy cows or pathogen exposure in poultry. We propose that genetic variations within these bivalent regions may capture key genotype-by-environment (GxE) interactions that are often missed by neutral markers. This hypothesis is strongly supported by recent findings in cattle, where integrating functional annotations significantly enhanced genomic prediction for climate resilience traits like heat tolerance [11]. Therefore, the strategy of prioritizing bivalent SNPs could be particularly valuable for developing robust breeding lines in livestock that maintain high performance across diverse rearing environments. Importantly, the predictive utility of epigenetically prioritized SNPs is likely to be trait- and tissue-dependent, as functional elements identified in the most relevant tissues (e.g., muscle for growth, adipose for fat deposition) have been shown to yield the greatest benefit in genomic prediction models [6,10]. Consistent with this view, our ongoing work on pigmentation traits in fish indicates that skin-derived regulatory annotations outperform those from other tissues, underscoring the importance of context-specific epigenomic prioritization.

Overall, this epigenome-informed prioritization strategy offers a practical route toward designing low-density, functionally enriched SNP panels that reduce genotyping costs while maintaining prediction accuracy across generations and populations. As functional genomic resources continue to expand in livestock species, integrating chromatin-state annotations into GS frameworks may provide an effective and scalable solution for improving breeding efficiency in large-scale animal agriculture.

### Limitations and future directions

Although this study generated high-resolution histone modification maps and demonstrated their application to GS, several aspects remain open for future improvement. In this study, we prioritized four histone marks (H3K4me1, H3K4me3, H3K27me3, and H3K27ac) and one adult tissue (muscle), as these modifications are well-established indicators of promoter and enhancer activity and are closely linked to growth regulation. Although this design provides a robust foundation for functional annotation, it may not fully capture the broader regulatory complexity across diverse tissues and developmental stages. Expanding to additional histone marks (e.g., H3K36me3, H3K9me3) and other functional layers such as ATAC-seq or Hi-C would further refine functional annotations. Second, while our prioritized SNPs showed strong predictive and cross-population performance, experimental validation of their causal roles will be an important next step. Finally, the general applicability of this strategy to other traits and aquaculture species remains to be systematically assessed.

Future work should therefore focus on integrating multi-omics datasets, extending analyses to diverse tissues and developmental stages, and testing the transferability of this approach across species and traits. The SNP sets identified here already provide practical resources for the development of optimized genotyping panels, and their validation in breeding programs will further accelerate the transition toward functionally informed GS in aquaculture.

## 5. Conclusions

In summary, this study establishes the first chromatin state–guided framework for SNP prioritization in GS of shrimp. By integrating multiple histone modification signals, we identified functional SNP subsets that improve predictive accuracy, reduce dependence on high-density genotyping, and enhance cross-population stability. The strong performance of E6 SNPs underscores the central role of bivalent promoter/enhancer regions in growth regulation and environmental adaptability. This work provides a theoretical and methodological basis for building “epigenetically informed” marker systems, advancing molecular breeding in aquaculture from traditional structural markers toward functional, biologically grounded markers. Future work should focus on validating these prioritized SNP panels in real-world breeding programs.

## List of abbreviations

CREs: Cis-regulatory elements
CUT&Tag: Cleavage under targets and tagmentation
CV: Cross-validation
FAANG: Functional annotation of animal genomes consortium
GBLUP: Genomic best linear unbiased prediction
GEBV: Genomic estimated breeding value
GO: Gene Ontology
GRM: Genomic relationship matrix
GS: Genomic selection
GWAS: Genome-wide association study
LD: Linkage disequilibrium
LMM: Linear mixed model
MAF: Minor allele frequency
PCA: Principal component analysis
PTMs: post-translational modifications
SNP: Single nucleotide polymorphism
TSS: Transcription start site
VCF: Variant Call Format
WGS: Whole-genome sequencing

## Ethics declarations

### Ethics approval and consent to participate

Protocols for animal use in this study were approved by the Institutional Animal Care and Use Committee (IACUC) of Ocean University of China.

### Consent for publication

Not applicable.

### Data availability

The resequencing datasets analyzed in this study were generated as part of ongoing selective breeding programs in our laboratory and are not publicly available due to their proprietary nature. These data may be made available from the corresponding author upon reasonable request. The CUT&Tag datasets used in this study include two categories: (i) seven embryonic developmental stages (blastula, gastrula, limb bud embryo, larva in membrane, nauplius I, nauplius III, and nauplius VI), which are publicly available from our previous publication (NCBI BioProject accession: **PRJNA1224423**) [33]; and (ii) newly generated adult muscle tissue data, which have been deposited in Figshare (https://figshare.com/) with a private link (https://figshare.com/s/a9b187bdd1f6fe2cc523). In addition, building upon these datasets, we generated CUT&Tag data for four later larval developmental stages—zoea I, zoea III, mysis I, and mysis III. These newly generated datasets have also been submitted to the NCBI BioProject database under accession number PRJNA1304883 (access link: https://dataview.ncbi.nlm.nih.gov/object/PRJNA1304883?reviewer=pan86d4b9sq8g6d9qkieplrnbs) All newly generated datasets will be made publicly available upon publication of this article.

## Competing interests

The authors declare no competing interests.

## Funding

This work was funded by the Project of Sanya Yazhou Bay Science and Technology City (SCKJ-JYRC-2023-62), National Natural Science Foundation of China (32200667), and Shandong Provincial Special Funds for Taishan Scholars (tsqn202306104).

## Author contributions statement

J.S. and Z.Y. conceived and designed the study. J.S. performed the methodology development, investigation, formal analysis, data curation, software implementation, validation, and prepared the original draft. Z.L., M.S., and M.Y. contributed to methodology and investigation. M.M. and D.Z. assisted with visualization and manuscript revision. Z.B. provided supervision, project administration, and resources. Q.Z. and J.H. contributed resources and participated in manuscript editing. Z.Y. supervised the project, acquired funding, and led the writing, review, and editing process. All authors reviewed and approved the final manuscript.

## Acknowledgements

We acknowledge the support of the High-Performance Biological Supercomputing Center at the Ocean University of China for this research.

## Declaration of generative AI and AI-assisted technologies in the writing process

During the preparation of this work the authors used ChatGPT-5.2 (OpenAI) for assistance in language polishing. After using this tool, the authors reviewed and edited the content as needed and take full responsibility for the content of the publication.

## Notes

### Competing Interest Statement

The authors have declared no competing interest.

