## Supplementary figures for "Epigenome-informed prioritization of bivalent chromatin SNPs enhances genomic prediction robustness: a proof-of-concept study in Pacific white shrimp (*Litopenaeus vannamei*)"

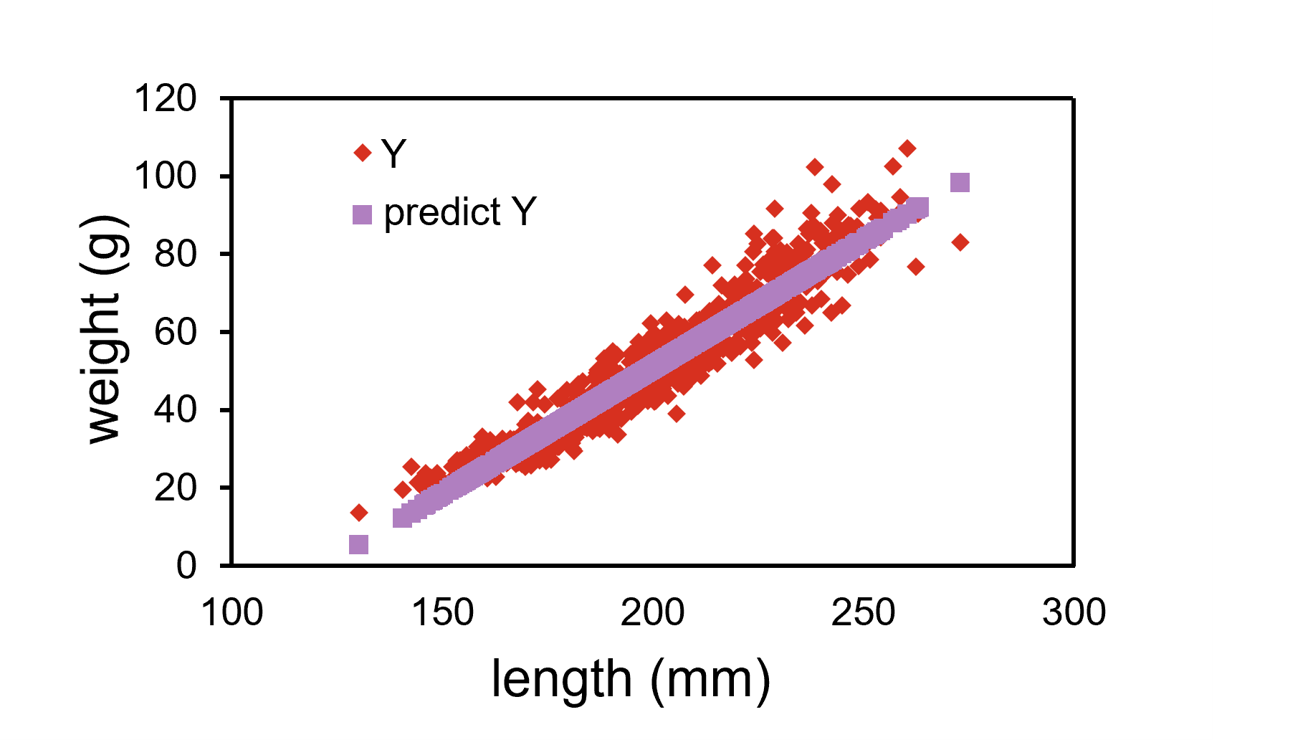

**Fig. S1.** Linear regression of body length on body weight. Red dots represent observed values, and the purple line indicates the fitted regression. A strong positive correlation was observed (R² = 0.918).

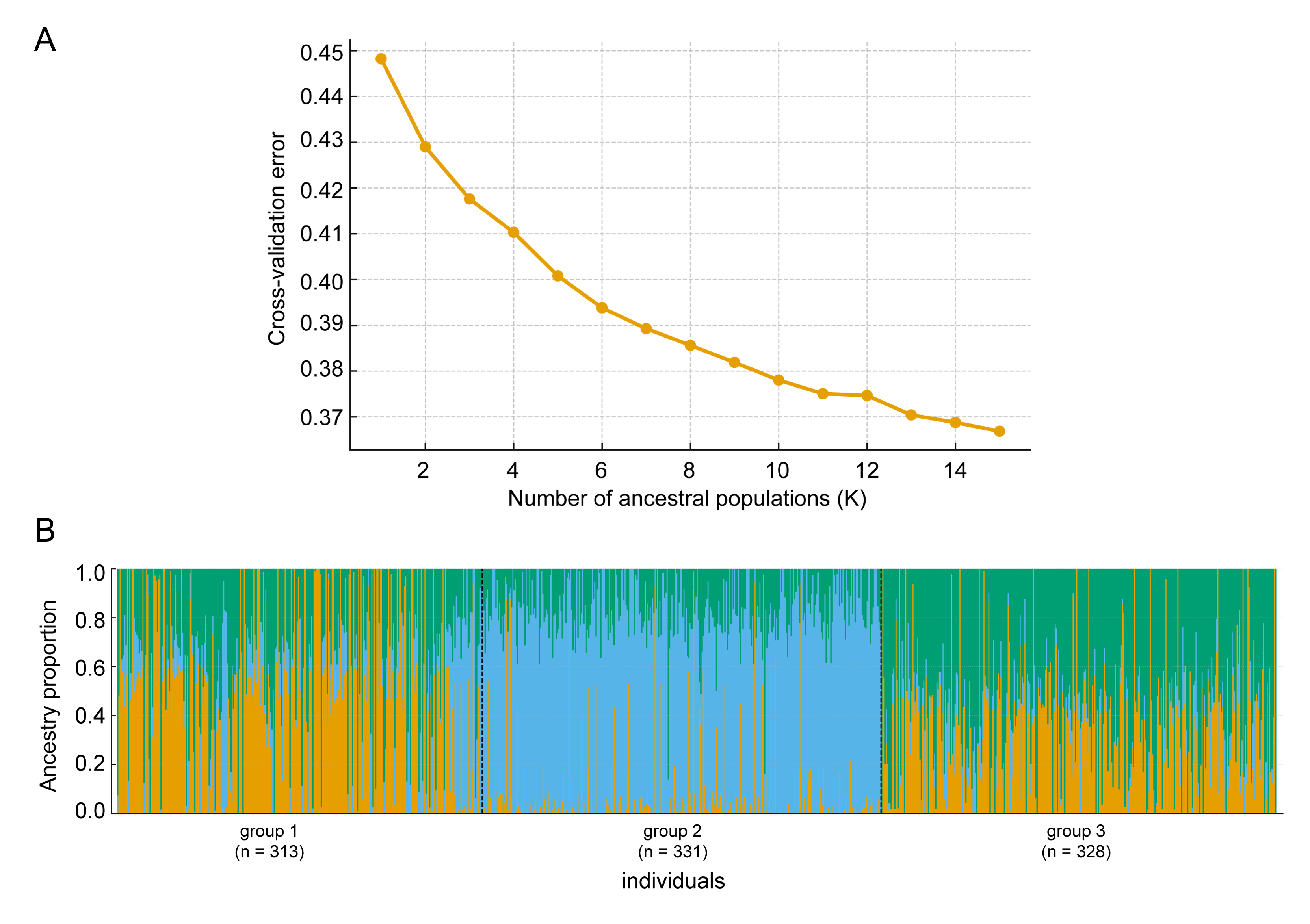
**Fig. S2.** Model-based population structure analysis of *Litopenaeus vannamei* populations inferred using ADMIXTURE. (A) Cross-validation (CV) error across different numbers of ancestral populations (K = 2–15). The CV error decreases gradually with increasing K, without showing a clear minimum. (B) Ancestry proportion bar plot for K = 3. Each vertical bar represents an individual, and different colors denote the estimated ancestry proportions from distinct genetic components. Individuals are grouped according to their sampling populations (group 1: n = 313; group 2: n = 331; group 3: n = 328), showing clear genetic differentiation and varying degrees of admixture among groups.

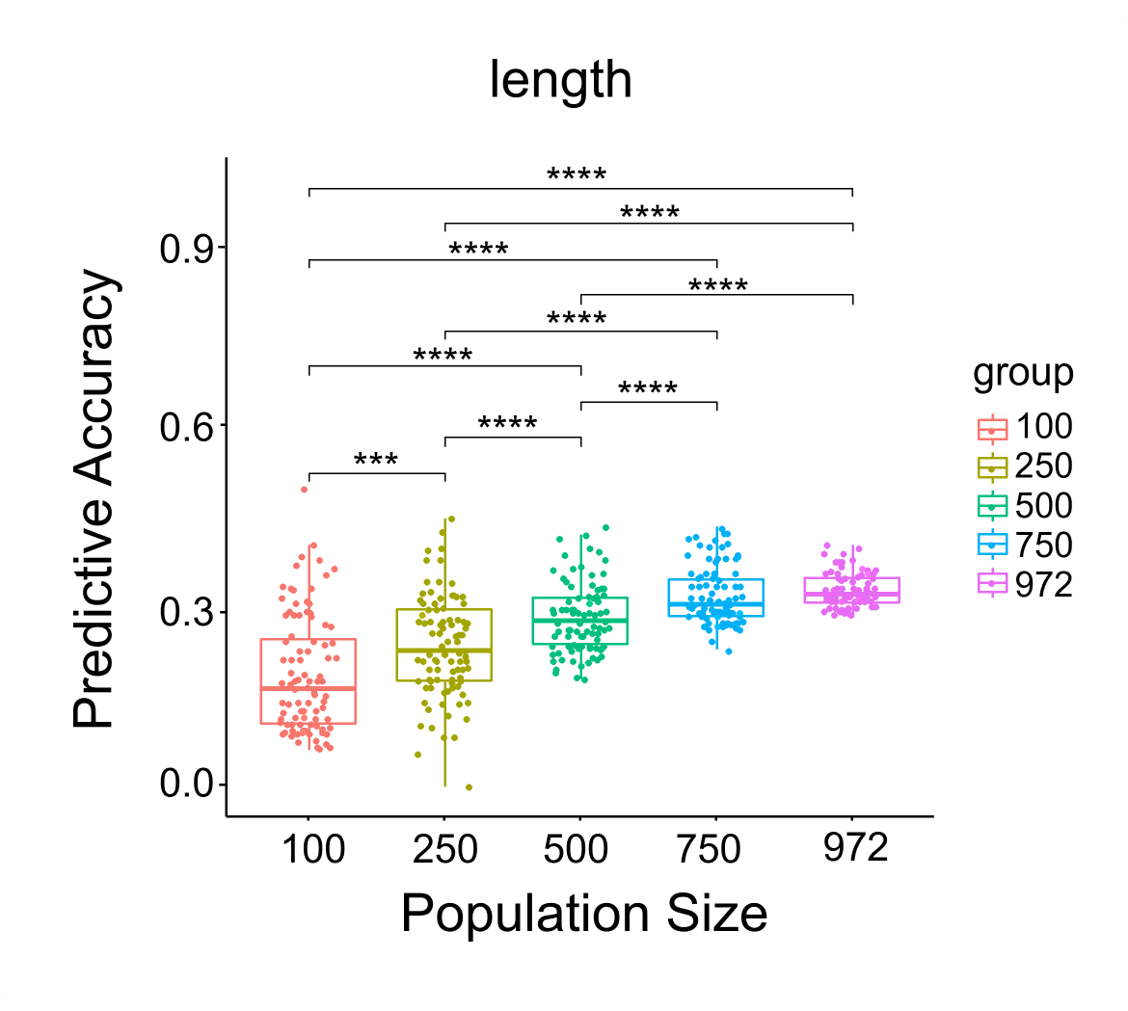

**Fig. S3.** Effect of training population size on prediction accuracy at a fixed density of 10k SNPs, evaluated using BayesA. For each size, SNPs were randomly sampled five times and assessed using 20 repetitions of cross-validation, withholding 20% of individuals for validation set in each iteration. Accuracy improved with larger training sets (*** *p* < 0.001, **** *p* < 0.0001) and plateaued beyond ~750 individuals.

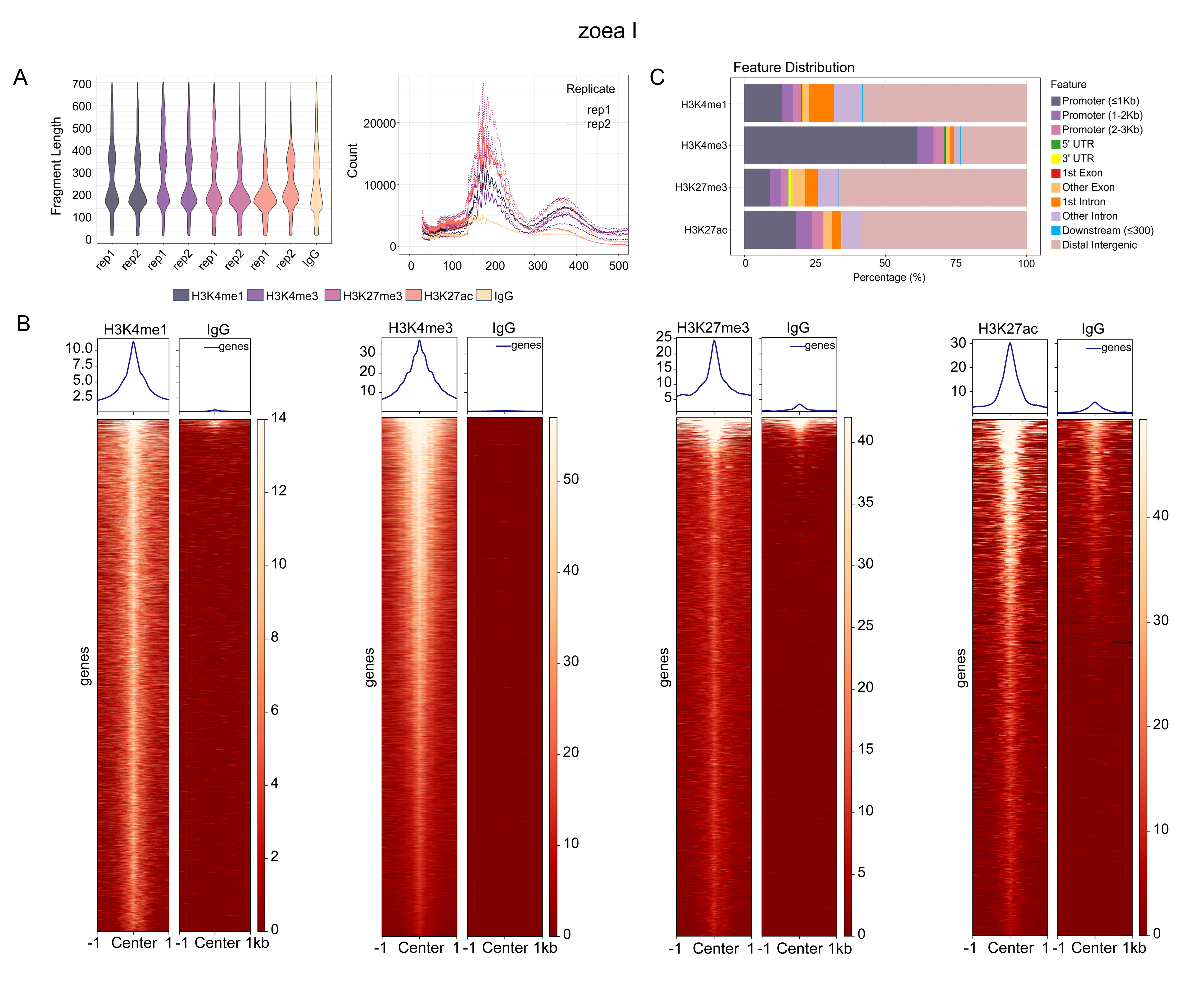

**Fig. S4.** Quality assessment of CUT&Tag data at the zoea I stage. (A) Fragment size distribution. Plots showing the proportion (left) and raw counts (right) of DNA fragments across different sizes. Histone modification samples display strong enrichment of mononucleosome-sized fragments, whereas the IgG control samples exhibit a random fragment size distribution. (B) Peak signal enrichment. Heatmaps (bottom) and line plots (top) showing CUT&Tag signal enrichment within ±1 kb of peak summits for the four histone modifications (H3K4me1, H3K4me3, H3K27me3, and H3K27ac), compared with the random signal distribution observed in the IgG control. (C) Distribution of peaks across genomic features.

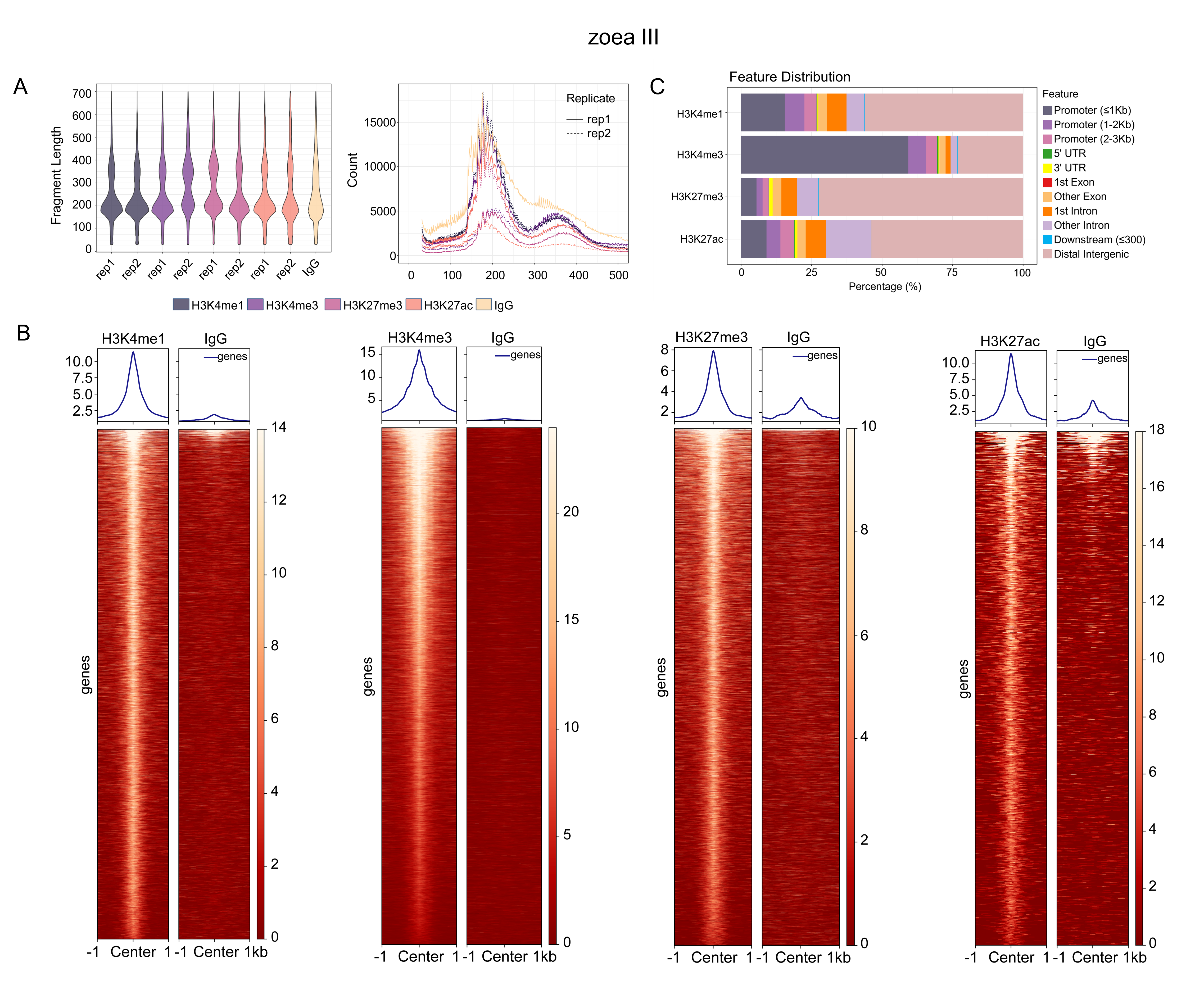
**Fig. S5.** Quality assessment of CUT&Tag data at the zoea Ⅲ stage. (A) Fragment size distribution. Plots showing the proportion (left) and raw counts (right) of DNA fragments across different sizes. Histone modification samples display strong enrichment of mononucleosome-sized fragments, whereas the IgG control samples exhibit a random fragment size distribution. (B) Peak signal enrichment. Heatmaps (bottom) and line plots (top) showing CUT&Tag signal enrichment within ±1 kb of peak summits for the four histone modifications (H3K4me1, H3K4me3, H3K27me3, and H3K27ac), compared with the random signal distribution observed in the IgG control. (C) Distribution of peaks across genomic features.

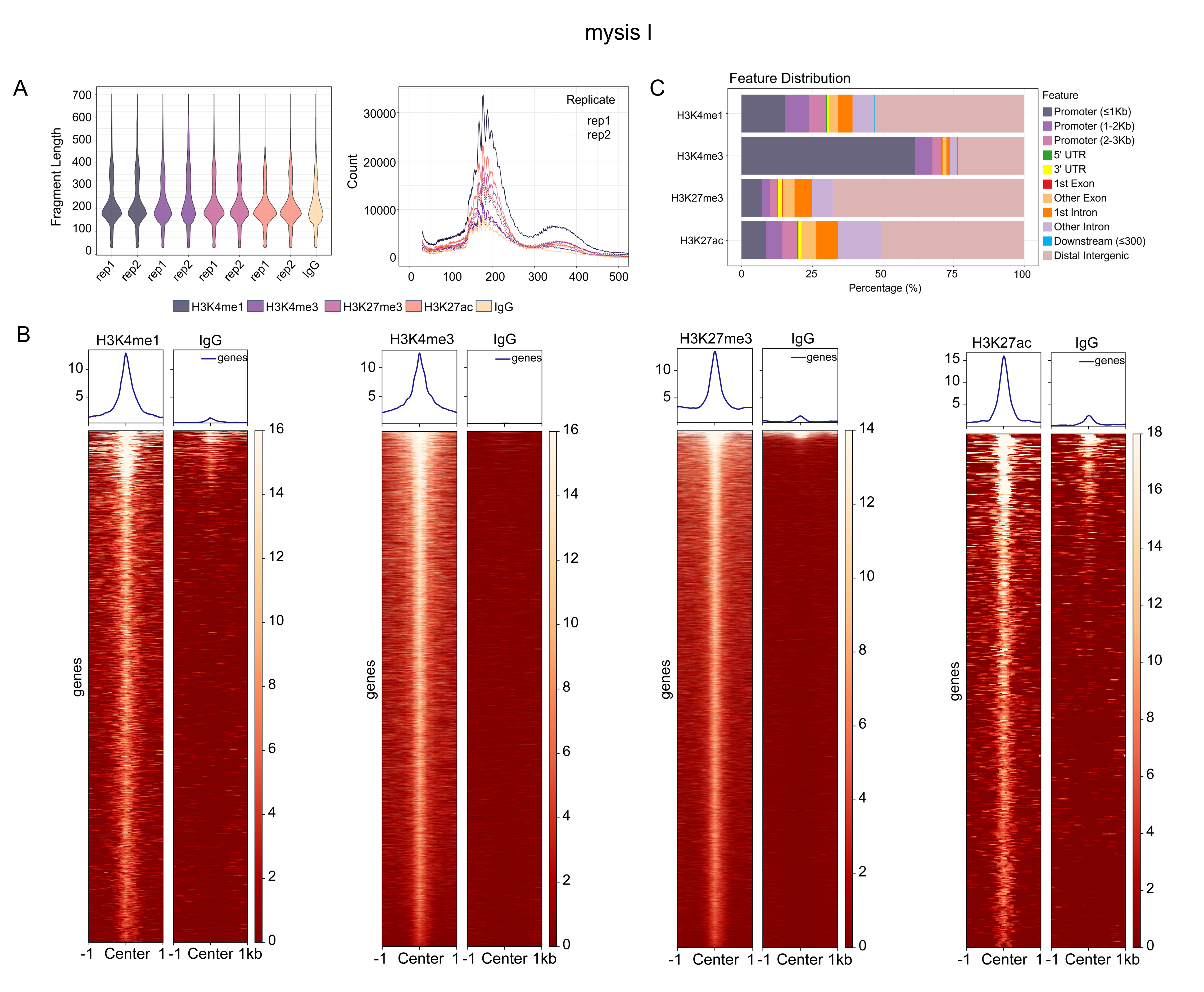

**Fig. S6.** Quality assessment of CUT&Tag data at the mysis Ⅰ stage. (A) Fragment size distribution. Plots showing the proportion (left) and raw counts (right) of DNA fragments across different sizes. Histone modification samples display strong enrichment of mononucleosome-sized fragments, whereas the IgG control samples exhibit a random fragment size distribution. (B) Peak signal enrichment. Heatmaps (bottom) and line plots (top) showing CUT&Tag signal enrichment within ±1 kb of peak summits for the four histone modifications (H3K4me1, H3K4me3, H3K27me3, and H3K27ac), compared with the random signal distribution observed in the IgG control. (C) Distribution of peaks across genomic features.

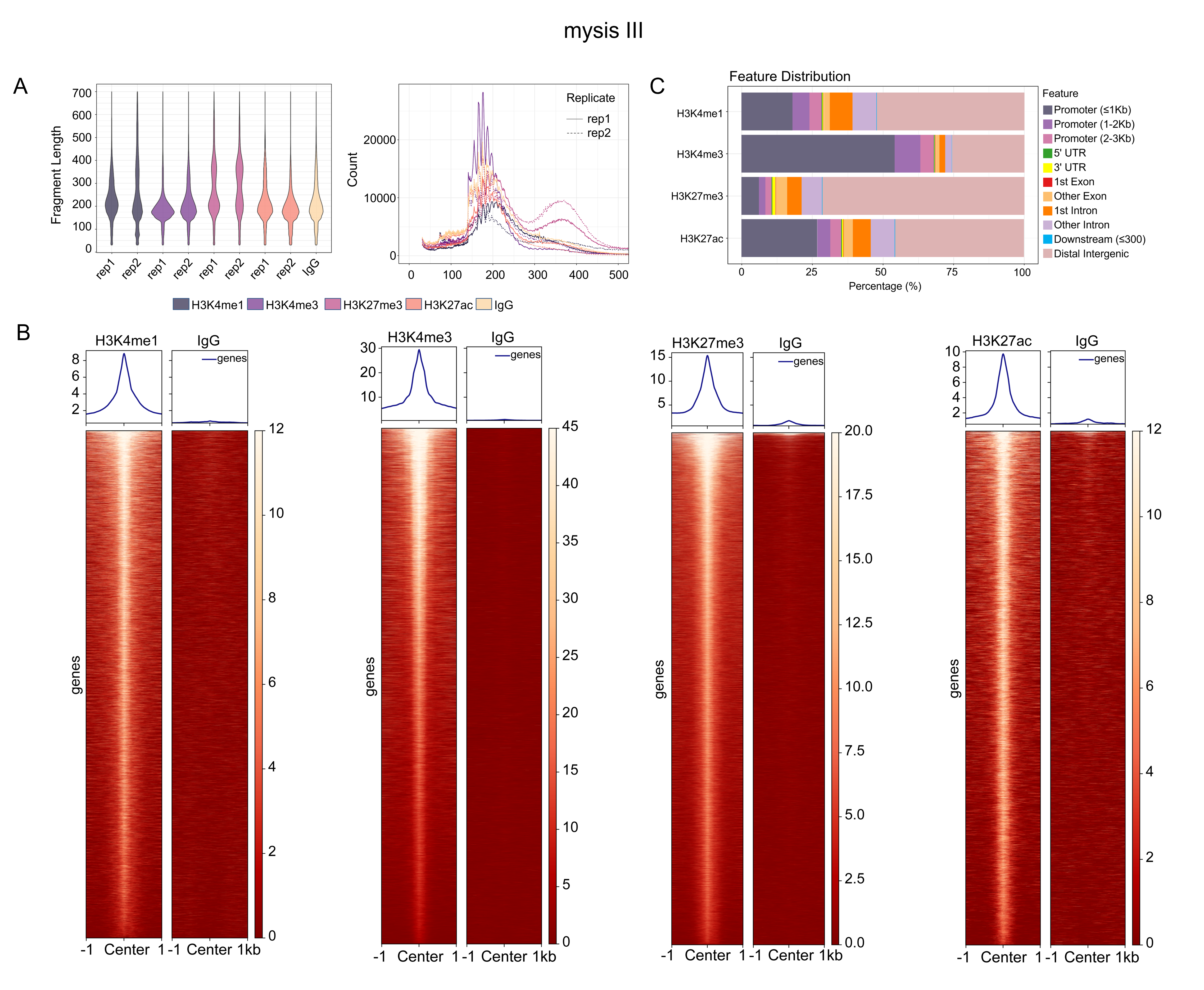

**Fig. S7.** Quality assessment of CUT&Tag data at the mysis Ⅲ stage. (A) Fragment size distribution. Plots showing the proportion (left) and raw counts (right) of DNA fragments across different sizes. Histone modification samples display strong enrichment of mononucleosome-sized fragments, whereas the IgG control samples exhibit a random fragment size distribution. (B) Peak signal enrichment. Heatmaps (bottom) and line plots (top) showing CUT&Tag signal enrichment within ±1 kb of peak summits for the four histone modifications (H3K4me1, H3K4me3, H3K27me3, and H3K27ac), compared with the random signal distribution observed in the IgG control. (C) Distribution of peaks across genomic features.

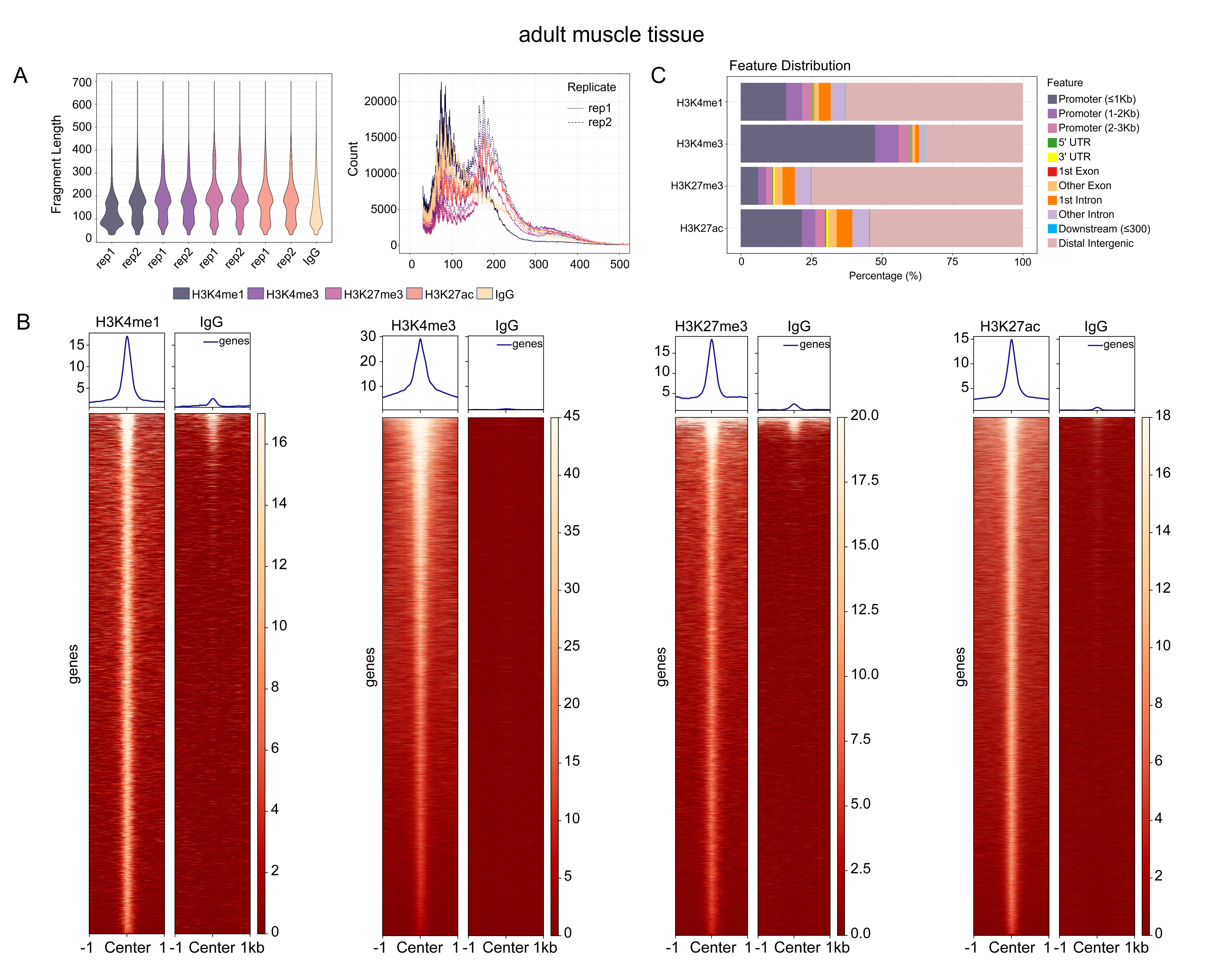

**Fig. S8.** Quality assessment of CUT&Tag data at the adult muscle tissue. (A) Fragment size distribution. Plots showing the proportion (left) and raw counts (right) of DNA fragments across different sizes. Histone modification samples display strong enrichment of mononucleosome-sized fragments, whereas the IgG control samples exhibit a random fragment size distribution. (B) Peak signal enrichment. Heatmaps (bottom) and line plots (top) showing CUT&Tag signal enrichment within ±1 kb of peak summits for the four histone modifications (H3K4me1, H3K4me3, H3K27me3, and H3K27ac), compared with the random signal distribution observed in the IgG control. (C) Distribution of peaks across genomic features.

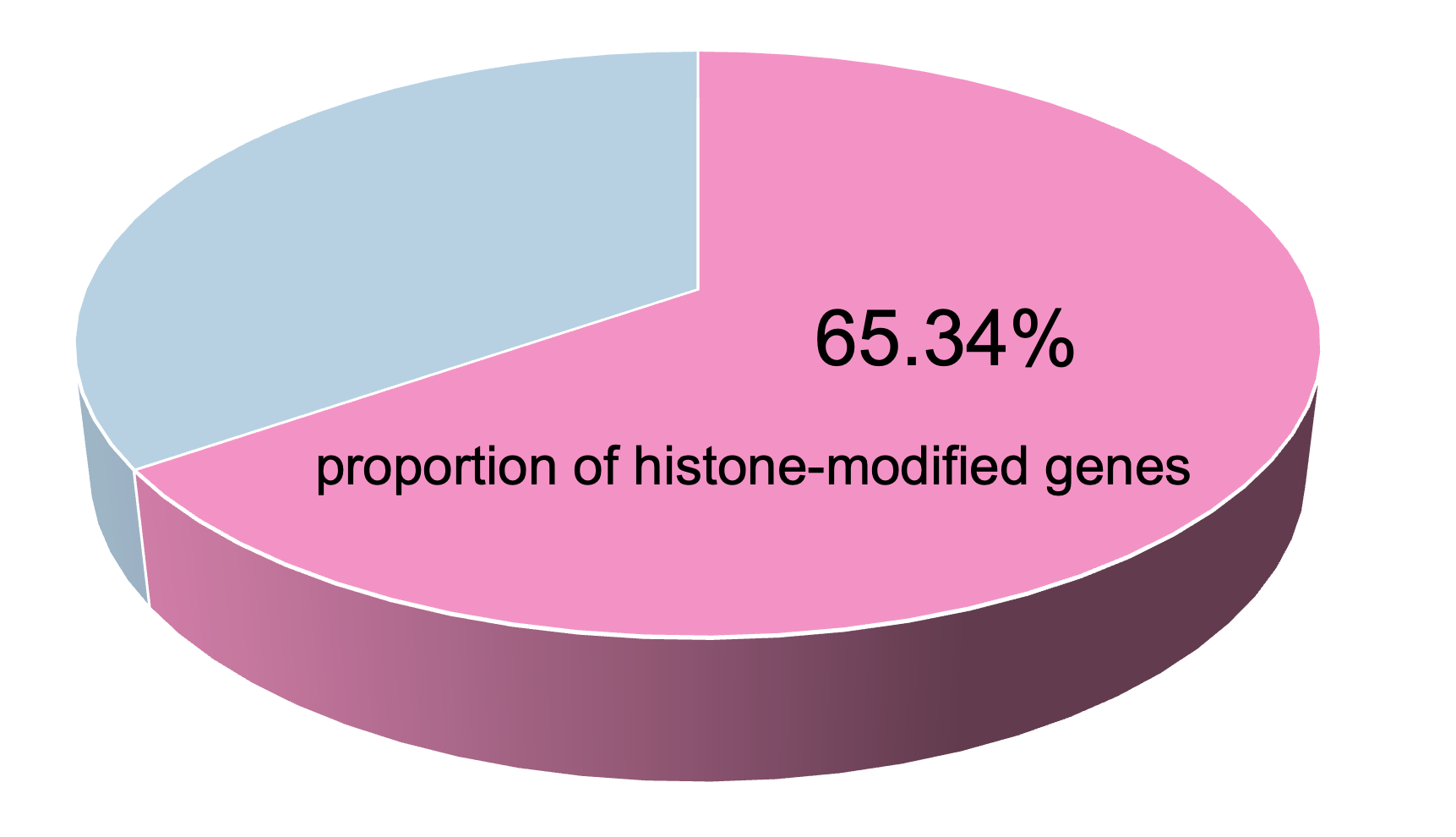

**Fig. S9.** Gene annotation of CUT&Tag peaks during embryonic development of *Litopenaeus vannamei*. CUT&Tag peaks from embryonic stages overlapped with 65.34% of annotated genes in the genome, indicating widespread association of histone modifications with coding regions and suggesting abundant cis-regulatory information during development.

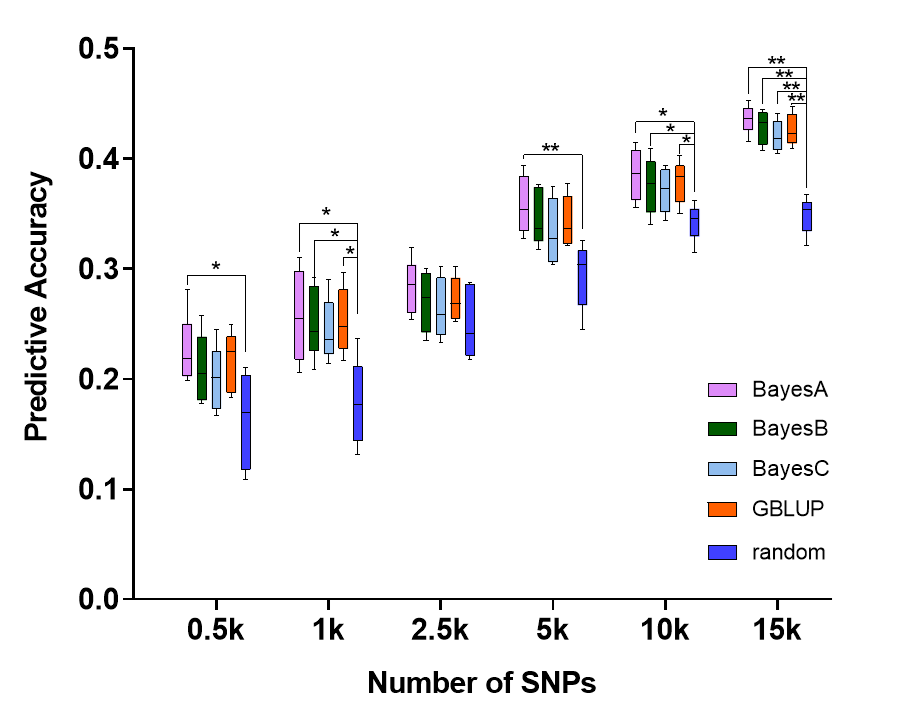

**Fig. S10.** Validation of SNPs from the muscle bivalent promoter/enhancer state (E6) across different genomic selection (GS) models. Four GS models (BayesA, BayesB, BayesC, and GBLUP) were used to evaluate the predictive accuracy of SNPs located in the muscle E6 chromatin state for body length across five SNP densities (0.5k–15k). For each density, SNPs were randomly sampled five times and assessed using fivefold cross-validation repeated five times. Across all models, SNPs from the E6 state consistently outperformed random SNPs. Significance levels: **p* < 0.05, ***p* < 0.01.
